# Interactions between natural selection and recombination shape the genomic landscape of introgression

**DOI:** 10.1101/2021.10.12.462572

**Authors:** Maud Duranton, John Pool

**Affiliations:** Laboratory of Genetics, University of Wisconsin-Madison, Madison, WI, USA

## Abstract

Hybridization between lineages that have not reached complete reproductive isolation appears more and more like a common phenomenon. Indeed, speciation genomics studies have now extensively shown that many species’ genomes have hybrid ancestry. However, genomic patterns of introgression are often heterogeneous across the genome. In many organisms, a positive correlation between introgression levels and recombination rate has been observed. It is usually explained by the purging of deleterious introgressed material due to incompatibilities. However, the opposite relationship was observed in a North American population of *Drosophila melanogaster* with admixed European and African ancestry. In order to explore how directional and epistatic selection can impact the relationship between introgression and recombination, we performed forward simulations of whole *D. melanogaster* genomes reflecting the North American population’s history. Our results revealed that the simplest models of positive selection often yield negative correlations between introgression and recombination such as the one observed in *D. melanogaster*. We also confirmed that incompatibilities tend to produce positive introgression-recombination correlations. And yet, we identify parameter space under each model where the predicted correlation is reversed. These findings deepen our understanding of the evolutionary forces that may shape patterns of ancestry across genomes, and they strengthen the foundation for future studies aimed at estimating genome-wide parameters of selection in admixed populations.

## Introduction

Speciation has long been seen as a gradual process during which two populations evolve barriers to gene flow until reproductive isolation is complete (Orr 1995; Coyne and Orr 2004). Now, many speciation genomic studies have shown that it is more of a dynamic process that can involve more than two populations (Novikova *et al*. 2016; Van Belleghem *et al*. 2021) and where hybridization is a common phenomenon (Payseur and Rieseberg 2016) that might also sometimes favor speciation (Jacobsen and Omland 2011; Schumer *et al*. 2015, 2018a; b; Blanckaert and Bank 2018). There is indeed a grey zone of speciation where diverging lineages can still exchange genes (Roux *et al*. 2016). Therefore, many species genomes are in fact a mosaic made of different ancestry. Introgression, that is the exchange of genetic material between diverging populations through hybridization, can have different effects on the receiving genome. In most cases, if introgression has a fitness consequence at all, it is probably deleterious either because introgressed DNA fragments can carry deleterious variants (especially if the donor population is small and thus has a high genetic load) (Bierne *et al*. 2002) or because they carry locally adaptive alleles not fit for the receiving population’s environment (Arnegard *et al*. 2014). The presence of Bateson-Dobzhansky-Muller incompatibilities (BDMIs) (Bateson 1909; Dobzhansky 1937; Muller 1942), in which an allele from one population at one locus has poor fitness when combined with an allele from a second population at a second locus, can also make introgression deleterious. Indeed, BDMIs are thought to be a major mechanism that contribute to the buildup of reproductive isolation and are expected to naturally arise when populations are diverging without gene flow (Masly and Presgraves 2007; Presgraves 2010). Nonetheless, incoming gene flow can also be beneficial through adaptive introgression. This process allows linked beneficial alleles that have already been tested by natural selection to enter a new genetic background (Fisher 1937; Racimo *et al*. 2017; Martin and Jiggins 2017). There are now several examples of species that have acquired adaptive phenotypes through introgression such as mimicry in *Heliconius* butterflies (Dasmahapatra *et al*. 2012), seasonal camouflage in the snowshoe hares (Jones *et al*. 2018), and altitude adaptation (Huerta-Sánchez *et al*. 2014) and malaria resistance in humans.

A consequence of selection that acts on introgressed alleles is that patterns of introgression are often heterogenous along the genome (Martin and Jiggins 2017). Understanding how hybrid genomes and patterns of introgression are modulated by natural selection is a major goal in evolutionary biology. To do so, it is necessary to take into account variations of recombination rate along the genome, as recombination can modulate the strength of selection through linkage disequilibrium (Nachman and Payseur 2012; Harris and Nielsen 2016; Juric *et al*. 2016). In the absence of selection, no consistent relationship between recombination rate and introgression level is expected. In contrast, for many species a positive correlation has been observed between introgression and recombination, such as in human with Neanderthal ancestry (Sankararaman *et al*. 2016; Juric *et al*. 2016; Schumer *et al*. 2018b), in hybrid populations of swordtail fishes (Schumer *et al*. 2018b), within populations of Heliconius butterfly (Martin *et al*. 2019) and between subspecies of house mice (Janoušek *et al*. 2015). These positives correlations are predicted by theory when introgression is deleterious. In highly recombining regions, neutral introgressed alleles can more easily recombine away from deleterious ones and persist in the genome (Barton and Bengtsson 1986). Whereas in low recombining regions where linkage disequilibrium is higher, deleterious variants when removed bring with them more neutral ones (Charlesworth *et al*. 1993), which leads to greater reductions in introgression levels.

A contrasting relationship between recombination rate and introgression level was observed for the *Drosophila* Genetic Reference Panel (DGRP). Like most North American populations of *Drosophila melanogaster*, this North Carolina population resulted from an admixture of populations likely originating from Europe and Africa (David and Capy 1988; Caracristi and Schlötterer 2003; Duchen *et al*. 2013; Kao *et al*. 2015; Bergland *et al*. 2016). This admixture was estimated to have occurred about 1,600 generations ago, based on the lengths of ancestry tracts (Pool 2015; Corbett-Detig and Nielsen 2017). Within the DGRP sample of 205 sequenced inbred strain genomes, a strong negative correlation has been observed between the minor African ancestry and recombination (Pool 2015; Corbett-Detig and Nielsen 2017), in contrast to the predictions of both neutral and BDMI models. This pattern serves as a motivating observation for the present study.

To our knowledge, the only published model predicting a negative correlation between introgression and recombination rate is based on simulations in which diverging populations experience substantial genetic load and each one may fix a distinct set of deleterious variants during an isolation of 2*N* generations (Kim et al. 2018). In that study, negative correlations were observed in models involving recessive deleterious variants (generating heterosis in admixed populations) and/or introgression flowing from a larger population into a smaller one. However, it is not clear that such a model should be responsible for the pattern observed in the DGRP population, in which admixture occurred between large source populations that only diverged on the order of 0.01*N* generations ago (Sprengelmeyer et al. 2020).

Here, we hypothesize that positive selection in admixed populations can also generate negative correlations between recombination rate and introgression level, even if both populations contribute the same number of favored variants. The key reason is that favored variants from the minor population will traverse a greater frequency change, and thus have a greater impact on ancestry levels (compared to major population favored variants), and this imbalance will be greater in low recombination regions where random neutral sites are more likely to be linked to selected sites. As a conceptual example, suppose minor and major populations initially admix in 10% vs. 90% proportions. And simplistically, assume that a random neutral site has a 50% chance of being linked to a selected variant if it’s in a low recombination region, but only a 10% chance of linkage to a selected variant if it’s in a high recombination region. If selection is fully successful in favoring beneficial variants, then the expected minor population ancestry proportion in low recombination regions is: (50% of sites not linked to selection × 10% minor population ancestry) + (25% of sites linked to favored major population variant × 0% minor population ancestry) + (25% of sites linked to favored minor population variant × 100% minor population ancestry) = 30%. Whereas, the expected minor population ancestry for high recombination regions is: (90% of sites not linked to selection × 10% minor population ancestry) + (5% of sites linked to favored major population variant × 0% minor population ancestry) + (5% of sites linked to favored minor population variant × 100% minor population ancestry) = 14%. This predicted excess of minor population ancestry in low recombination regions due to positive selection is qualitatively similar to the genetic load model (Kim et al. 2018) but in contrast to expectations from incompatibility models (Martin and Jiggins 2017).

In this study, we use simulation to identify the relationships between recombination and introgression generated by positive selection in an admixed population. We also perform similar simulation analyses using models of pairwise incompatibilities, in part to test whether any specific BDMI scenarios depart from the general expectation of a positive recombination-introgression correlation. We simulated whole genomes of *D. melanogaster* within two source populations that admix in different proportions to generate a third one that evolves on its own for 1,600 generations. After this time, we analyzed patterns of introgression along the genome and the relationship between the minor population ancestry and recombination rate. For the model of directional selection, we varied the number of favored variants present in the population, the fraction of variants presents in each population, the average selection coefficient of the variants, and the admixture proportion. For the model with pairwise incompatibilities, we varied the average fitness impact of the incompatibilities, the dominance of the variants, and the admixture proportions. We show that models with only positive variants often yield negative correlations between minor ancestry and recombination (particularly in cases where the two source populations contribute equal numbers of favored variants) but that depending on the parameters, positive correlations can also be generated. Furthermore, we also show that spatial correlation of population ancestry level along the genome may help distinguish between models of fewer strong versus many weak favored variants. We also confirmed that incompatibilities tend to produce positive introgression-recombination correlations, but we discovered that this correlation can also be reversed in scenarios where one population carries dominant incompatibility partner alleles and the other carries recessive partner alleles. These results will guide the interpretation of evolutionary genomic data from admixing/introgressing taxa, while also improving prospects for the development of novel inference methods to estimate evolutionary parameters from such cases.

## Results

### Directional selection - General patterns

We simulated a model where two populations (P1 and P2) containing each a certain fraction of the total number of positively selected variants admix in different proportions (*p*) to form a third one (P3). Test simulations confirmed that we could scale down the population size to 10,000 with no meaningful influence on average genomic ancestry landscapes (Supplementary Figure 1). Sex ratio was also found not to have any notable impact on these results, and was otherwise fixed at 50% females/males.

We then varied the average selection coefficient of the beneficial variants 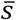, along with the total number of favored variants (*n*) and the fraction of favored variants present in each population (*f*) (Figure 1A and 2A). In scenarios with equal numbers of favored variants (*i*.*e*., same fitness, *f* = 50%-50%) and unequal admixture proportions from each source population (*p* = 10%-90% or 30%-70%), we found that negative correlations between recombination rate and minor population ancestry were consistently produced (Figure 2B, C, F and G yellow, Supplementary Table1 and Supplementary Figures 2 and 3), in line with the predictions described in the Introduction (the rise of minor population alleles has a relatively greater impact on ancestry compared to favored major population alleles, and this effect is greatest in regions of low recombination).

**Figure 1.**
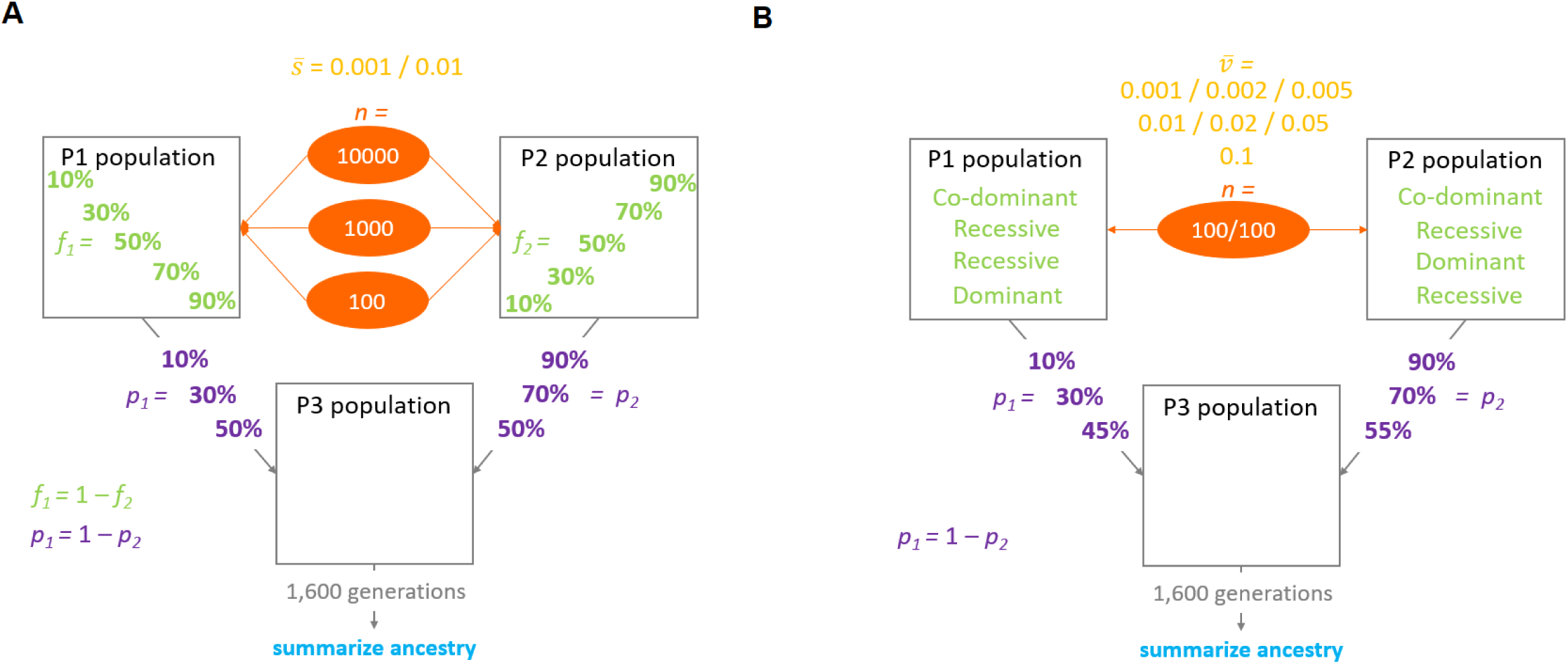
Schematic representation of the models used in the simulations. Two source populations P1 and P2 are mixed in different proportions (*p*, admixture proportion) to form a third population P3 that evolves on its own. After 1,600 generations population, P3 ancestry is analyzed to determine which one of P1 or P2 is the minority ancestry at the end of the simulations. **A**. Model of directional selection where a certain number of positively selected variants (*n*, number of variants) is distributed between P1 and P2 populations in different proportions (*f*, fraction of beneficial variants). Variants’ selection coefficients are drawn from an exponential distribution with a predetermined mean value 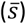. **B**. Model of pairwise incompatibility where the P1 population contains 100 fixed alleles each involved in a pairwise negative interaction with one of 100 alleles fixed within the P2 population. Each negative interaction reduces the fitness of the individual who carries it by a certain value (*ν*) drawn from an exponential distribution with a predetermined mean value 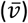 Three different models of dominance were simulated: one with only co-dominant alleles, one with only recessive alleles, and one with one population containing only dominant alleles and the other only recessive ones.

**Figure 2:**
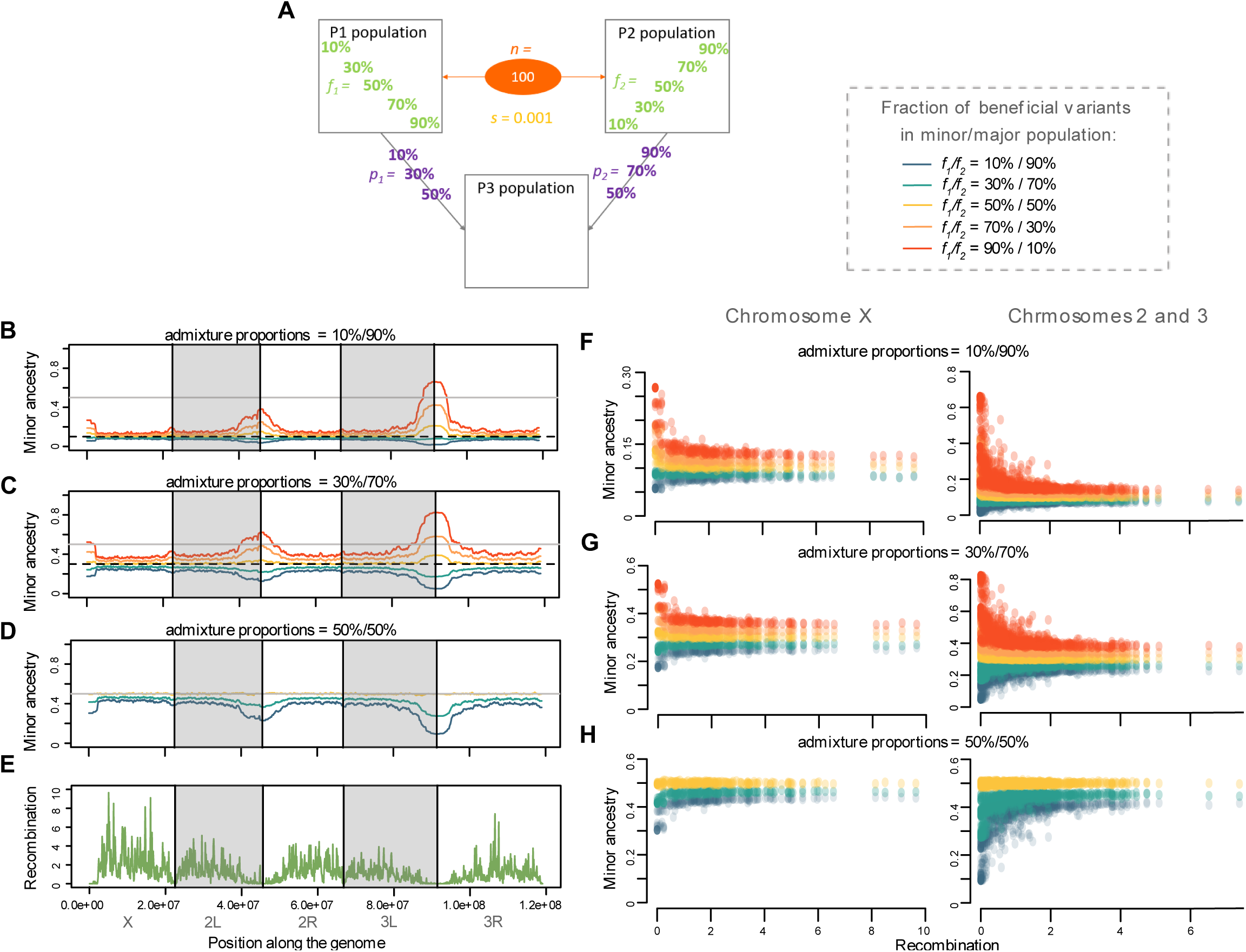
Directional selection can produce negative or positive correlations between recombination rate and minor population ancestry. **A**. Depiction of the specific parameter space of directional selection presented in the following panels, in terms of admixture proportions (*p*) and beneficial variant fractions (*f*) for cases with 100 selected variants (*n*) with average 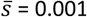. The left graphs depict the average minor ancestry proportion (across 1,000 replicates) along the genome when *p*_*1*_ and *p*_*2*_ are: **B**. 10% and 90%, **C**. 30% and 70%, and **D**. 50% and 50%. **E**. Recombination rate (sex-averaged crossover frequency in cM/Mb) along the genome of *D. melanogaster*. The right panels depict the relationship between minor ancestry proportion and recombination for the X chromosome and the autosomes when *p*_*1*_ and *p*_*2*_ are: **F**. 10% and 90%, **G**. 30% and 70%, and **H**. 50% and 50%. Colors in panels B-D and F-H represent the fraction of beneficial variants contributed by each source population, as indicated on the right of panel A.

In contrast, scenarios in which the admixing source populations contributed unequal numbers of beneficial variants produced more diverse correlations between recombination and minor population ancestry. Positive correlations were observed in some scenarios if the ancestry that was minor at the end of the simulations was predominantly disfavored over time as proposed by Martin and Jiggins (2017). This pattern was often observed if an initially-major (or co-equally contributing) source population had a higher fitness (*i*.*e*. contributed most of the beneficial variants; Figure 2F, G, H light and dark blue; Supplementary Figures 2 and 3 when *f* = 10%-90% / 30%-70%), or if the minor population contributed most favored variants but selection was pervasive enough to change it into the major ancestry by the time of sampling (Supplementary Figure 2C when *f* = 90%-10% / 70%-30%). This switch in major/minor ancestries between the beginning and the end of the simulation was more likely to happen if selection was strong, and in regions of low recombination. Due to the autosomes’ lower recombination rate than the X chromosome in the *D. melanogaster* genome, ancestry reversals happened relatively more frequently on the autosomes (See Supplementary Table 1 and Supplementary Figure 2 and 3 when *p* = 90%-10% / 70%-30% and *f* = 10%-90% / 30%-70%). These results are all consistent with the general expectation that positively selected variants will bring more linked neutral variation with them in regions of low recombination (Maynard Smith and Haigh 1974). In addition, models of asymmetric fitness between admixing source populations may lead to synergistic frequency increases between beneficial variants from the more-fit population, particularly in regions of lower recombination. In light of both of these factors, the more-fit population’s ancestry should increase more rapidly in low recombining regions, potentially generating a positive correlation between the minor ancestry proportion and recombination.

Nonetheless we found some particular cases where even if the initially-major population had a higher number of beneficial variants, there was a negative correlation between minor ancestry proportion and recombination, especially in cases with fewer but stronger favored variants. This is true for 3 models on the X and the autosomes: when n = 100, s = 0.01, and (A) *p* = 10-90% and *f* = 10%-90% (B) *p* = 10-90% and *f* = 30%-70% and (C) *p* = 30-70% and *f* = 30%-70%, see Supplementary Figure 2B and 3B) and for 2 other models only on the X chromosome (when n = 1000, s = 0.01, and (A) *p* = 10%-90% and *f* = 30*%-70% and* (B) *p =* 30%-70% *and f =* 30%-70%, see Supplementary Figure 2D). These results may reflect the competition between processes favoring positive correlations between recombination and minor population ancestry (previous paragraph) and the unequal magnitudes of beneficial frequency changes that favor negative correlations (Introduction).

Negative correlations between recombination and minor population ancestry were also observed if the initially-minor population contributed more beneficial alleles but did not rise to become the major ancestry at the time of sampling (See Supplementary Figure 2A and 3A when *p* = 10%-90% / 30%-70% and *f* = 90%-10% / 70%-30%). Here, both differences in population fitness and the unequal magnitudes of beneficial frequency changes lead the minor population ancestry to increase, particularly in regions of low recombination, thus generating negative correlations. In summary, the correlations between recombination rate and minor population ancestry generated by positive selection in admixing populations are complex. Negative correlations are observed when major and minor source populations contribute equal numbers of beneficial variants, and also when the minor population contributes more beneficial variants but remains the minor ancestry. In contrast, positive correlations are usually generated if a major (or equally-admixing) population contributes more beneficial variants, or if a favored minor population becomes the major population.

### Number versus strength of selected sites

The above results suggest that the correlation between recombination rate and minor population ancestry holds information about the nature of selection in admixed populations. Such information may be useful in extending the evolutionary parameter inferences that are possible from population genomic data. However, some parameters may have superficially similar influences on population ancestry in high versus low recombination regions, such as the number versus strength of selected sites, and hence the inference of these parameters could be somewhat confounded. Our simulations offer the possibility to compare models with the same genome wide average level of selection 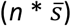 with different number of positively selected variants (Figure 3 and Supplementary Table 2).

**Figure 3:**
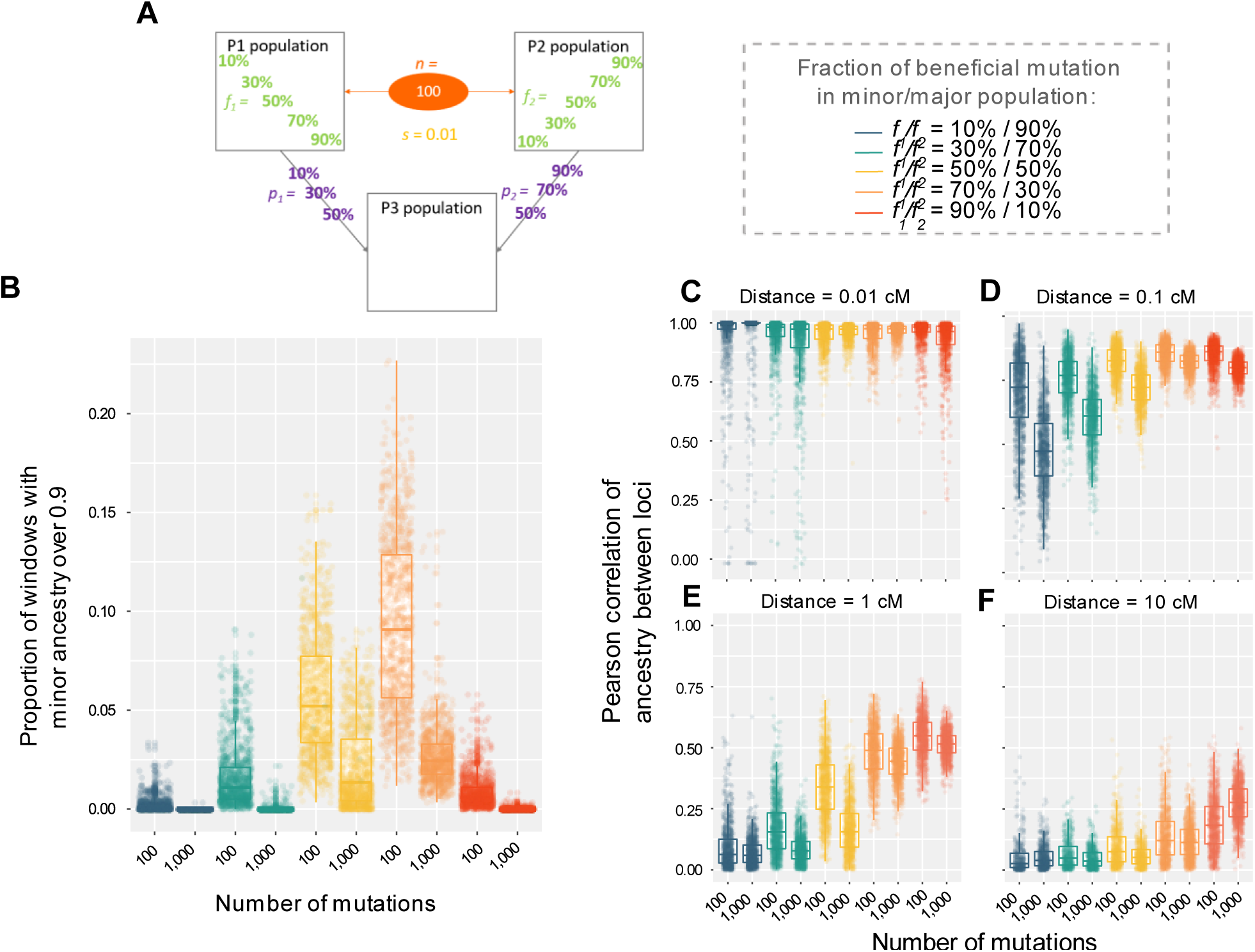
Differences between models of directional selection with the same level of selection genome-wide but different numbers of selected variants. **A**. Depiction of the specific parameter space of directional selection presented in the following panels: models with either *n* = 100 variants and an average 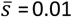, or *n* = 1,000 variants and an average 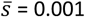, were examined for different beneficial variant fractions (*f*) for cases in which *p*_*1*_ = 10% and *p*_*2*_ = 90%. **B**. Distribution among replicates of the proportion of genomic windows with minor ancestry above 0.9. **C**. Spatial correlations of ancestry along chromosomes, comparing between the same models as described above. Each point represents a replicate and measures of Pearson correlation involving all 100k b windows separated by a distance of: **D**. 0.01 cM, **E**. 0.1 cM, **F**. 1cM and **G**. 10cM. Colors in panels D-G represent the fraction of beneficial variants contributed by each source population, as indicated on the right of panel A.

Here, we compared patterns of p_2_ ancestry along the genome for two models with an average summed selection coefficient of 1 (either n = 100 and 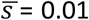 or n = 1,000 and 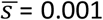), since models with more pervasive selection (*i*.*e*. summed selection coefficient of 10) often resulted in complete fixation of one ancestry (Supplementary Table 2). We can see that population ancestry patterns tend to be smoother when there is a smaller number of variants with on average a higher selection coefficient compared to a case with a higher number of variants with a low selection coefficient (Figure 3B). When we focus more precisely on the spatial correlation of population ancestry along the genome, we can see that at a short genetic distance, 0.01 cM, there is a very high spatial correlation for both models and we cannot distinguish the two kinds of model (Figure 2D and Supplementary Table 2). In the same way, at a very high genetic distance, 10cM, spatial correlations are very low and there are almost no differences between the models (Figure 3H and Supplementary Table 2). However, in between these two distances there is a clear tendency for spatial correlations to be higher for models with a lower compared to higher number of positively selected variants (Figure 3F and G and Supplementary Table 2). It also seems that for both kind of models, correlations tend to be stronger as the proportion of p2 is higher. Therefore, the genetic distance at which it is easier to distinguish between few versus many variants seems to depend on the admixture proportion, meaning that future inferential studies may wish to consider spatial correlations of population ancestry along chromsosomes at multiple scales of genetic distance.

### Pairwise incompatibilities

In our second model, both source populations contain 100 fixed variants that are involved in pairwise negative epistatic interactions with the 100 variants fixed in the other population (Figure 1B). As for the directional selection model, we confirmed that population size could be scaled down to 10,000 and that skewed sex ratios had little effect on ancestry patterns (Supplementary Figure 4).

We then varied the average fitness impact of the incompatibility 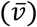as well as the admixture proportion of the two populations (*p*) and the dominance of the variants involved in the incompatibilities (*d*) (Figure 4A). As expected, we showed that in most cases there is a positive correlation between the minor ancestry proportion and recombination (Figure 4, Supplementary Table 3 and Supplementary Figure 5 and 6). Indeed, neutral introgressed variants are more likely to recombine from incompatible loci in highly recombining regions and therefore to persist there. However, for one model in particular – when the major population only carries recessive variants and the minor one only dominant variants – the relationship between the minor ancestry proportion and recombination is sometimes negative (Figure 4E and J, Supplementary Table 3 and Supplementary Figure 5C and 6C). For example, on the autosomes, when the dominant population starts at 30% and the average fitness effect is between 0.001 and 0.01, there is a negative relationship (Figure 4E and J). When the fitness effect is stronger, the minor population at the end of the simulation is no longer the dominant population but the recessive one, and in that case the relationship is positive (Supplementary Figure 5C). In the same way for the X chromosome, where recombination is higher, when the proportion of the dominant population is 30% and the average fitness effect is at least equal to 0.005, there is a negative relationship between recombination and minor ancestry proportion (Supplementary Figure 6C). Similarly, we find that there is also a range of parameter space generating negative correlations when the initial admixture proportion of the dominant population is 45% (Supplementary Figure 5C and 6C).

**Figure 4:**
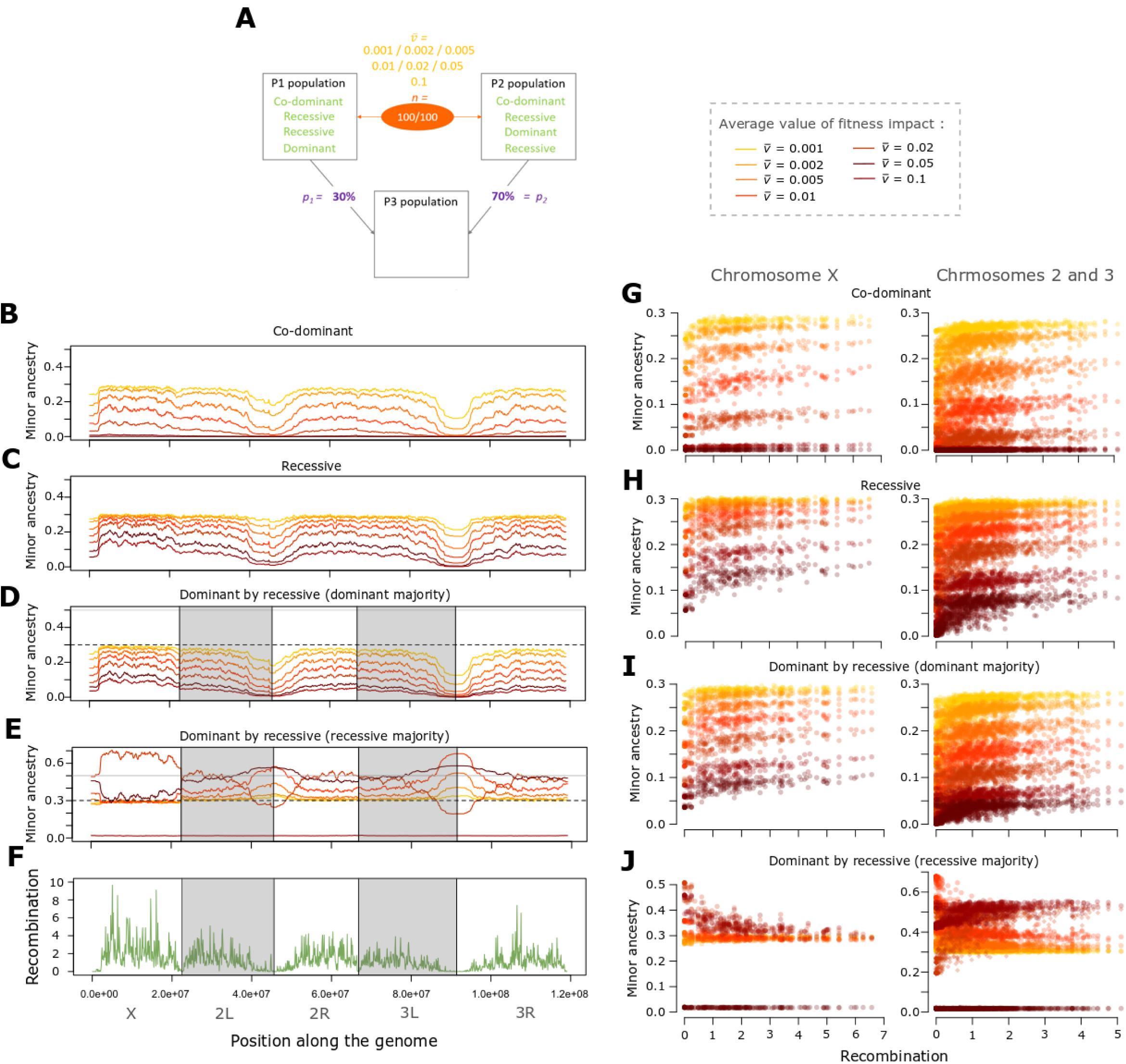
Incompatibilities usually generate positive correlations between recombination rate and minor population ancestry. **A**. Depiction of the specific parameter space of the incompatibility presented in the following panels, for an admixture model in which *p*_*1*_ = 30% and *p*_*2*_ = 70%, and there are 100 pairs of incompatible variants (*n*). Results are shown for several mean fitness effects 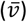 and dominance scenarios. The left graphs depict the average minor ancestry proportion (across 1,000 replicates) along the genome when incompatibility fitness effects are: **B**. co-dominant, **C**. recessive, **D**. dominant in the major population and recessive in the minor, or **E**. recessive in the minor population and dominant in the major. **F**. Recombination rate (sex-averaged crossover frequency in cM/Mb) along the genome of *D. melanogaster*. The right panels depict the relationship between minor ancestry proportion and recombination for the X chromosome and the autosomes when variants are: **G**. co-dominant, **H**. recessive, **I**. dominant in the major population and recessive in the minor, or **J**. recessive in the major population and dominant in the minor. Colors in panels B-D and F-H represent the mean fitness effect of incompatibilities, as indicated on the right of panel A.

To try to understand why there is this negative relationship in this particular parameter space, we used two-locus analytical models to predict the frequency change of two autosomal loci involved in an incompatibility with one allele being dominant and the other one recessive. We fixed the fitness reduction due to the incompatibility to 0.01 whether the dominant allele is homozygous or heterozygous, and took into account the probability that either one or both partners may be on the X chromosome. We did these calculations for different proportions of admixture and determined after 1,600 generation which ancestry (the dominant or recessive one) had increased during this time (Figure 5). Indeed, similarly to the model of directional selection, there was a positive correlation between minor ancestry and recombination when the minor ancestry decreased over time, and a negative one when it increased (unless the major ancestry switched). We confirmed that (with a fitness reduction of 0.01) there is indeed a particular parameter space where the recessive population starts between 60% and 70% and is still the major population at the end of the simulations even if its ancestry tends to decrease. This means that during the 1,600 generations, the dominant ancestry tends to increase but not enough to become the major population, which explains why there is a negative relationship between the minor ancestry proportion and recombination. If the recessive ancestry starts in lower proportion (below 60%) then it is decreasing over time and remains the minor population explaining why there is a positive correlation between recombination and introgression (Figure 5). If the recessive ancestry starts in higher proportion (over 70%) then it is increasing over time, meaning that the minor ancestry proportion is decreasing and thus generates a positive correlation between the minor ancestry proportion and recombination (Figure 5).

**Figure 5:**
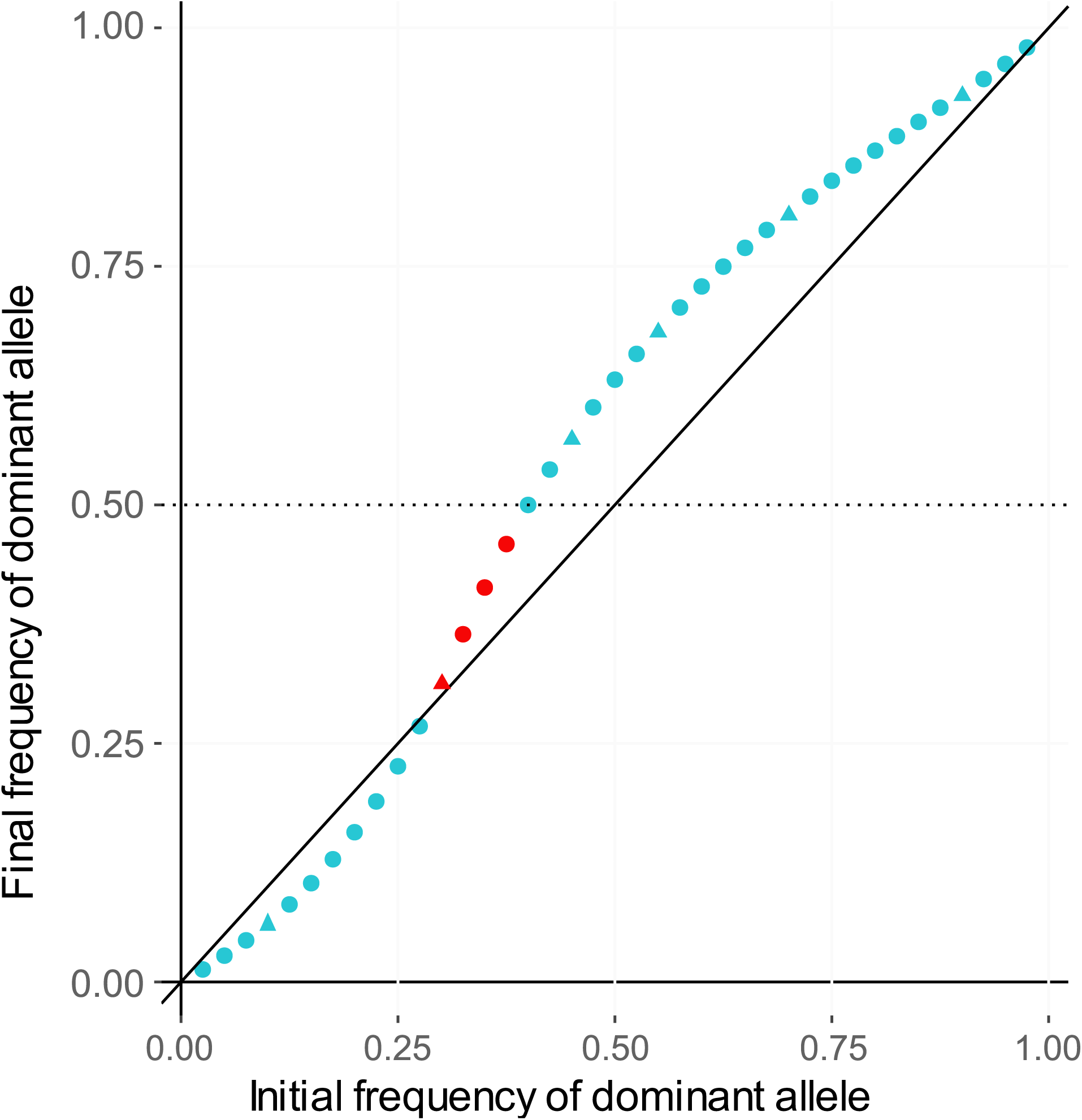
Theoretical expectation for the frequency change of incompatible variants under a dominant-by-recessive model. Results were calculated using a simple two-locus fitness model in which incompatible two-locus genotypes led to a relative fitness reduction of 0.01. Either or both partner loci could fall on the X chromosome with probability 20%, and were otherwise autosomal. No linkage between partner loci was modeled. In this plot, the x axis represents the initial admixture proportion of the dominant incompatibility partner locus, and the y axis is its predicted frequency after 1,600 generations of admixture. The black line of equation x=y indicates whether frequency has increased or decreased over time. Triangles represent frequencies that were included in our simulation study and red points indicate scenarios in which the predicted correlation between minor ancestry and recombination is negative.

These results can be explained by the interaction between the probability for an allele to be in a deleterious genotype and the absolute frequency shift that it undergoes. For example, in a model with the recessive ancestry starting at 10%, the dominant allele is going to be in a deleterious genotype 1% of the time whereas the recessive one is 10% of the time. Purifying selection is thus stronger on the recessive allele, and its (minority) ancestry will decrease more strongly in low recombination regions, leading to a positive correlation between introgression and recombination (Figure 5). However, when the recessive population starts at 70%, the recessive allele is deleterious 36% of the time and the dominant one is 49%, and yet overall the (minority) dominant ancestry is increasing, leading to a negative correlation. Here, the net increase of dominant ancestry may be attributable to the differing magnitudes of frequency shifts experienced by initially major versus minor alleles (as we suggested above for the directional selection model). In this case, the greater frequency reduction experienced by recessive incompatible alleles in this scenario may outweigh their lesser fitness consequences on average. Of course, there is also the possibility to have a switch of major/minor ancestry between the beginning and the end of the simulations. When the recessive population starts at 55%, the recessive allele is deleterious 38% of the time and the dominant one 30%. Here, the dominant ancestry increased enough to become the major ancestry, and there is a positive correlation between introgression and recombination (Figure 5).

## Discussion

In this paper we aimed at understanding how patterns of introgression along the genome are influenced by forms of positive and negative selection. We simulated two populations of *D. melanogaster* that are mixed in different proportion to form a third one. The source populations contained fixed variants that were either unconditionally positively selected or else involved in pairwise negative epistatic interaction with the variants fixed in the other source population. We showed that positive selection can sometimes generate negative correlations between the minor ancestry and recombination, and that incompatibilities generally result in positive correlations, but that for either model, the correlation can be reversed in some scenarios.

### Directional selection

In our model containing only unconditionally beneficial variants, we showed that when major and minor source populations contribute equal numbers of beneficial variants to an admixed population, negative correlations between recombination rate and minor ancestry proportion are generated (Supplementary Figures 3 and 4). We attribute this observation primarily to the greater ancestry frequency change entailed by fixing a minor population allele compared to a major population allele, and the greater impact of this imbalance on neutral variation in regions of low recombination.

In contrast, when there is a fitness difference between the two source populations (meaning here a difference in the fraction of variants present in the population *f*), we usually recover the expected relationship between minor ancestry and recombination (Supplementary Figure 3 and 4). The population with the higher fitness is more often under positive selection, and so its ancestry increases in frequency over time (particularly in low recombination regions), whether it started as the major or minor contributor. Indeed, when selection is strong enough there might be a switch between the minor ancestry at the beginning and the end of the simulations. Hence, either positive or negative correlations can be generated in cases of asymmetric fitness.

These results are comparable to what was observed in the presence of additive deleterious variants (Kim *et al*. 2018). When the receiving population has a lower fitness than the donor, introgressed alleles can be favored in low recombining regions (Kim *et al*. 2018), whereas this phenomenon is reversed if introgression is from a less fit into a more fit population. Thus, positive and negative correlations between minor ancestry and recombination can be generated by both the presence of deleterious and beneficial variants, depending on the relative fitness of the admixing population.

However, not all results can be explained by fitness differences between the two admixing populations. For example, when deleterious variants are recessive, an increase of introgression can be observed in low recombining regions when populations have the same fitness and even if the donor population as a lower fitness than the receiving one (Kim *et al*. 2018). This is explained by the fact that introgressed haplotypes generate associative over-dominance by preventing the expression of recessive deleterious variants (Ohta and Kimura 1970; Kim *et al*. 2018). As introgressed blocks are longer in low recombining regions, negative recombination-introgression correlations are produced. In contrast to the deleterious model, we find that a positive selection model can generate negative recombination-introgression correlations without fitness asymmetry or recessiveness. Although we have focused on additively beneficial variants, future studies may wish to examine the interplay between dominance and positive selection in admixing populations as well.

### Pairwise incompatibilities

In our model with source populations each containing 100 fixed variants involved in incompatibilities, we showed that when all variants are co-dominant or recessive, we always recover the expected correlation between minor ancestry and recombination, that is a positive one (Supplementary Figure 5A,D and 6A,D). Indeed, introgressed neutral variants can more easily recombined from those involved in incompatibilities and thus segregate in highly recombing regions. We also tested another model of dominance, where one population only contains dominant variants and the other only recessive variants. This particular scenario might happen in nature if one population went through a particular event of local adaptation and assuming that adaptation tended to favor either dominant or recessive variants. Indeed, alleles of local adaptation may sometimes be involved in genetic incompatibilities, especially if the two populations have been evolving in different environments. We find that in the particular parameter space in which the population containing the recessive variants starts as the major contributor, the correlation between minor ancestry and recombination can actually be negative (Figure 3E,J and Supplementary Figure 5 and 6).

To complement our simulation results, we also used mathematical predictions to estimate the expected frequency of each allele involved in the two-locus incompatibility for each of the 1,600 generations of our simulated admixture case to understand how this negative correlation is generated. We found that when the recessive population starts between 0.6 and 0.7, a negative correlation is indeed expected (Figure 5). Indeed, in this particular model of dominance there is an interaction between the probability of an allele to be deleterious (which determines the strength of purifying selection on it) and the frequency of this allele. In fact, what is really important is not the relative frequency reduction of each allele but the absolute one. An allele might have a smaller relative frequency shift as selection that acts on it is smaller but have overall a higher absolute frequency shift as it started at a higher frequency (analogous to the justification for our directional selection hypothesis). In this case, this allele ancestry is going to decrease faster even if selection is less strong. This correlation reversal parameter space is probably too particular to be widespread in nature. Nonetheless, the variability and magnitude and even direction of recombination-introgression correlations we observed under incompatibility models further underscores the complex task of making inferences about selection models in admixed populations.

### What about the particular case of D. melanogaster?

Qualitatively, we now know of three types of selection in admixed populations that are capable of generating a negative correlation observed between the minor ancestry proportion and recombination rate such as that observed in a North American DGRP population of *D. melanogaster* (Pool 2015). As previously demonstrated by Kim et al. (2018), an accumulation of deleterious variation between populations can lead to this outcome, especially if this variation is recessive and/or introgression is from a larger population into a smaller one. Importantly, the simulations of Kim et al. involved relatively smaller populations (*N*_*e*_ = 10,000) that had been separated for 2*N*_*e*_ generations before introgression occurred. Whereas in the case of our fly population, not only have effective population sizes been generally much larger (aside from a moderate out-of-Africa bottleneck), which should selection against deleterious variation, but also the divergence between the African and European source population of the DGRP only occurred within the past 0.01*N*_*e*_ generations (Sprengelmeyer *et al*. 2020). Hence, it is not clear that a sufficient quantity of deleterious variation should be expected to have fixed between these fly populations on such a time scale, in order for this model to generate a strong negative correlation.

Our study has identified two other ways in selection might generate a negative correlation. Regarding selection against deleterious incompatibilities, we identified a particular scenario involving consistent differences in the dominance of incompatibility alleles between source populations. Incompatibilities may well exist between the African and European source populations of the admixed DGRP population. European and African populations of *D. melanogaster* show signs of partial reproductive isolation (Wu *et al*. 1995; Yukilevich and True 2008; Lachance and True 2010; Kao *et al*. 2015), and a surprising abundance of ‘ancestry disequilibrium’ between unlinked loci in the DGRP population (Pool 2015) may point to the presence of epistatic selection of this nature. The dominance asymmetry required for this model to generate a negative correlation is conceivable – for example if incompatibilities were primarily due to adaptation in the European source population’s lineage, and this adaptation mainly involved recessive variants. However, it is not clear that incompatibilities should primarily be of such a type.

Finally, we show that across a somewhat broader parameter space, models of positive selection can generate negative correlations between recombination rate and minor population ancestry. In the context of the DGRP population, where it appears that admixture occurred between a majority European source population and a minority African source population, a negative correlation would probably be expected under positive selection under the simplest scenario in which these source populations contributed equal numbers of beneficial variants. The same prediction would also hold if a greater share of beneficial variants came from the African source population. Hence, the positive selection hypothesis may be the strongest candidate to explain the strong negative correlation observed in the DGRP fly population (where African ancestry ranges from 30% in low recombination regions to 10% in high recombination regions). However, we must emphasize that our study is not designed to identify a specific causative model, and that these different mechanisms are not mutually exclusive and they may all play a role in the observed relationship between recombination and ancestry.

## Conclusion

We have shown that positive selection in admixed populations can generate negative correlations between recombination and minor population ancestry, but that this correlation can be reversed if positive selection generally favors major population alleles or leads to a change in the major ancestry. These results are somewhat analogous to previous findings for models involving deleterious variation, which produces mainly negative correlations but with some exceptions. However as outlined above, the mechanisms underlying the predictions of these two directional selection models are somewhat different. We also confirm that positive correlations between recombination and introgression are generally expected from pairwise incompatibility models, but we find that here too, there are exceptions (specifically involving asymmetric dominance between source populations). Our results build on previous research to establish a more complete picture of the effects of selection on recombination-ancestry relationships (Figure 6).

**Figure 6:**
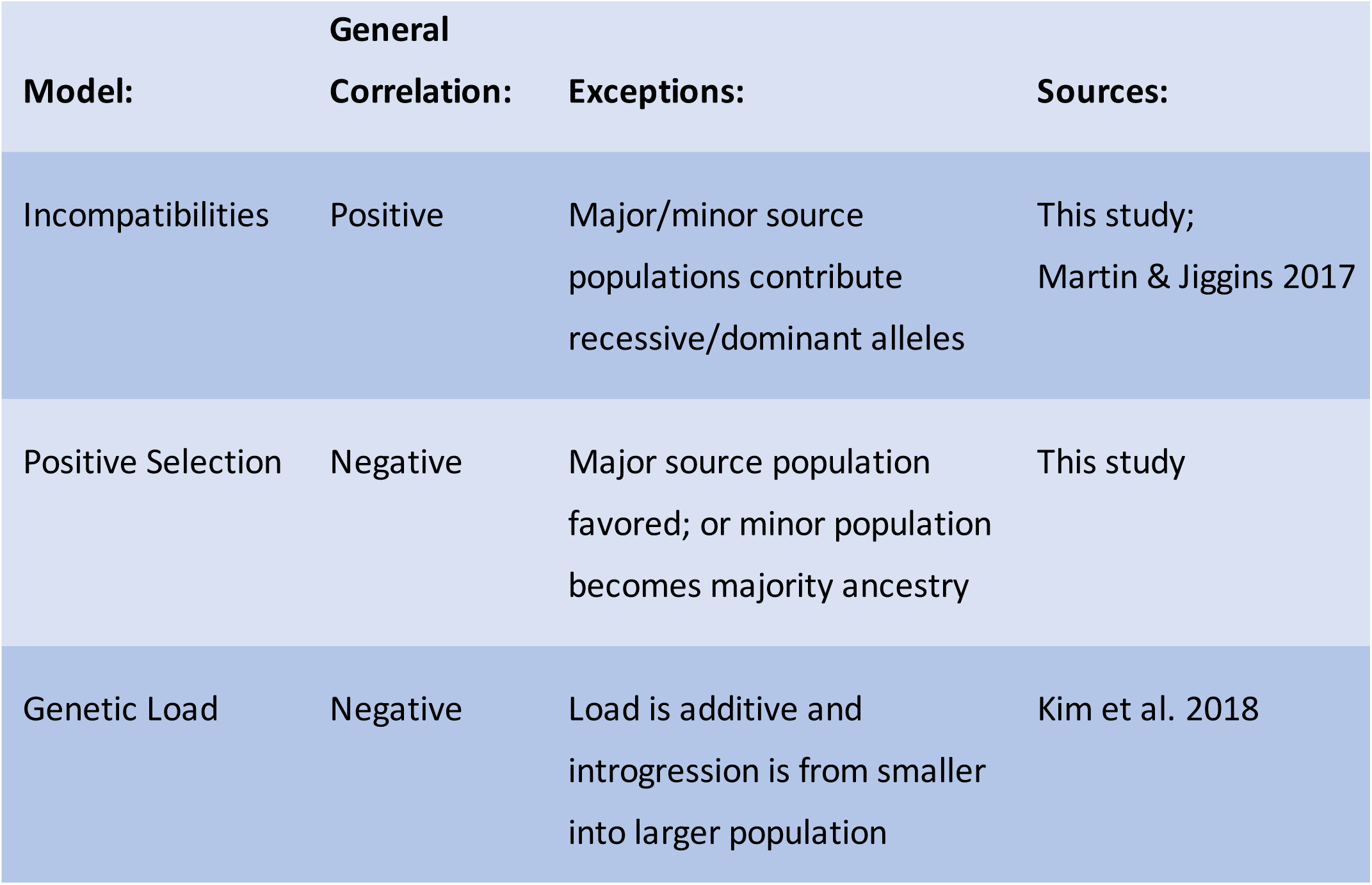
General relationships between minor population ancestry and recombination rate depending on the selection model, and known exceptions.

Given that any of the three models discussed above can generate either positive or negative correlations between recombination and ancestry, one obvious conclusion is that the direction of such a correlation is not sufficient to infer a causative selection model. In some cases, aspects of species biology and population history may inform the plausibility of different models (*e*.*g*. degree of reproductive isolation, effective population size, population genetic time-scale of isolation), as discussed above in the case of *D. melanogaster*. Otherwise, we suggest that future inferential studies will need to consider not only quantitative characteristics of this relationship, but also complementary aspects of genomic variation in admixed populations. Our analysis of spatial correlations in ancestry proportion (in the context of separating the number and strength of selected sites) provides just one example of such an approach. Further work is needed to identify additional features of ancestry variation in admixed populations that help to differentiate the above models. Nevertheless, the above findings bring new perspectives on how positive and negative selection interact with recombination to shape hybrid genomes and suggest promising directions for estimating the parameters involved.

## Materials and methods

### Basic structure of the model

To study the relationship between introgression and recombination, we simulated admixture between two populations using forward simulations in SLiM 3.5 (Haller and Messer 2019), assuming different selective effects. All files and scripts needed to run the simulations can be found at https://github.com/Mduranton/simulations_introgression_recombination. We wanted our model to reflect admixture between the European and African source populations of *Drosophila melanogaster* in North America, as represented by genomic data from the *Drosophila* Genetic Reference Population (Mackay *et al*. 2012). Therefore, we modeled the structure of *D. melanogaster* genome (119Mb in total), focusing on the two major autosomes (2 and 3) and the sex chromosome (X), using the recombination map estimated by (Comeron *et al*. 2012) with recombination only occurring within females. In the first generation before admixture, two populations with a sex-ratio of 0.5 are created. Then depending on the model, a certain number of fixed variants are randomly positioned along the genome of each population. As the X chromosome represents approximately 20% of the total genome size, on average 20% of variants occur on the X chromosome and the rest randomly fall along chromosome 2 and 3. Since all variants are fixed, the size of the two source populations does not matter; thus, to improve computational efficiency we choose to only start the simulations with 10 individuals in each population. In the second generation, a third population of 10,000 individuals with a sex-ratio of 0.5 is created from a certain fraction of population P1 and P2, depending on the model simulated. For computational feasibility, we choose to use only 10,000 individuals, but we also simulated different population sizes to confirm that this parameter has no noticeable influence on the relationship between introgression and recombination (see Supplementary Figure 1A,B,D,F and Supplementary Figure 4A,B,D,F). In the same way, as the actual sex ratio of *D. melanogaster* is not clearly identified, we simulated different sex-ratios to confirm that sex ratio does not strongly influence the relationship between recombination and introgression (see Supplementary Figure 1A,C,E,F and Supplementary Figure 4A,C,E,F). On the third generation, population P1 and P2 are removed. Finally, population P3 evolves on its own during 1,600 generations, which is approximately the time since admixture started between the European and African population of *D. melanogaster* in North America (Pool 2015; Corbett-Detig & Nielsen 2017). As we are only interested here in how the fixed variants present in the two first populations impact the relationship between introgression and recombination, the mutation rate was set to zero for all our simulations. We simulated 1,000 whole-genome replicates for each set of parameters.

In order to evaluate the level of introgression along the genome we tracked ancestry along the genome using a feature of SLiM, tree sequence files, that record genealogies all along the simulations (Haller *et al*. 2019). Tree sequence files were then analyzed using a custom python script and tools from pyslim v0.600 (Kelleher *et al*. 2018) and msprime v0.7.4 (Kelleher *et al*. 2016) that allow to trace the ancestry of different portions of the genome back to population P1 or P2 for each individual. In order to have the same effective population size along the whole genome, we only analyzed ancestry within females (the homogametic sex in *D. melanogaster*). Ancestry estimates were then average across individuals and replicates. We focused on the relationship between recombination and the minor ancestry proportion, meaning the minority ancestry at the end of the simulations, to reflect what can be observed in natural populations where the initial admixture proportions are not known.

### Directional selection

We created a model with only positively selected variants, with four different parameters (Figure 1A). The mean of the exponential distribution of selection coefficient, 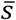, took two different values: 0.001 or 0.01. The number of variants present in the model, *n*, was either 100, 1,000, or 10,000. The proportion of variants fixed in population P1 and P2, *f*, was respectively either 10%-90%, 30%-70%, 50%-50%, 70%-30%, or 90%-10%. Lastly, *p*, the proportion of population P1 and P2 contributing to the creation of population P3, was respectively either 10%-90%, 30%-70%, or 50%-50%. When variants were (randomly) positioned along the genome, they were assigned a selection coefficient sampled from the exponential distribution previously defined. Depending on the parameters selected, the population contributing the most to population P3 could either have a higher or lower number of positive variants than the population that contributed the least. The combination of all the different parameters represents 78 different scenarios that were all replicated 1,000 times.

### Pairwise incompatibilities

In this model, populations contained fixed, otherwise-neutral variants that are involved in negative epistatic interactions and therefore deleterious when present with the interacting allele (Figure 1B). Incompatibilities were always generated between one allele fixed within population P1 and an allele present in population P2, in order to model Bateson–Dobzhansky–Muller incompatibilities (BDMI) (Dobzhansky 1937; Muller 1942; Coyne and Orr 2004). For example, population P1 is fixed for alleles “A” and “B” at the first and second locus respectively, whereas, population P2 is fixed for alleles “a” and “b”. Alleles “B” and “a” that have never been in the same genotype can prove to be deleterious when interacting. In our model, each population had 100 variants of this kind, generating 100 pairwise incompatibilities. As variants were randomly positioned along the genome, incompatible loci were not necessarily present on the same chromosome and incompatibilities could also be generated between an autosome and the X chromosome. There were three different parameters in our model. The average of the exponential distribution of the fitness reduction generated by the pairwise incompatibility, 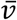, took 7 different values: 0.001, 0.002, 0.005, 0.01, 0.02, 0.05, or 0.1. The proportion of population P1 and P2 contributing to the creation of population P3, *p*, was either 10%-90%, 30%-70%, or 45%-55%. In this model population P1 was always the population that initially contributed the least to P3 population. Finally, *d* represents the dominance, as we simulated 3 different types of incompatibility: models where all incompatible alleles are co-dominant, or all recessive, or where one population contained only dominant alleles and the other only recessive ones. For the last model, the population containing recessive alleles could either be the major or minor contributor to P3 population. For each pairwise incompatibility, a value of fitness reduction (*ν*) was sampled from the exponential distribution previously defined. The fitness of each individual was then calculated depending on its genotype. For the co-dominant model, individuals that were homozygous at both incompatible loci (AAbb) had a fitness of 1-*ν*, individuals that were homozygous at one locus and heterozygous at the other (Aabb or AAbB) had a fitness of 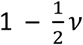, and individuals that were heterozygous at both loci (AaBb) had a fitness of 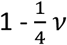. For the recessive model, only individuals that were homozygous at both loci (AAbb) had a fitness reduction equal to *ν*. Finally, for the dominant by recessive model, individuals that were homozygous with the incompatible allele at the recessive locus have a fitness reduction equal to *ν* if they presented one or two incompatible alleles at the dominant partner locus (AABb or AAbb if A is recessive and b dominant). The combination of all the different parameters represents 84 different scenarios that were all replicated 1,000 times.

We also generated mathematical predictions of the evolution of alleles frequency under the dominant-by-recessive model. To do so we calculated the expected frequency of two unliked variants one dominant and one recessive after 1,600 generations of admixture considering different initial frequencies and a fitness reduction of 0.01 for incompatible two-locus genotypes. Mirroring our simulations, we assume there is 20% chance that each variant is located on the X chromosome, and the sex-ratio is 0.5. We then estimated the frequency shift for each of four possibilities (both partners autosomal, recessive autosomal by dominant X-linked, dominant autosomal by recessive X-linked, and both X-linked) and then estimated an average final allele frequency weighted by the probability of each scenario.

### Analyses of ancestry

In order to determine if the correlation between minor ancestry proportion and recombination is positive or negative, we measured the correlation ratio between recombination rate and introgression for each scenario using the minor ancestry proportion average across the 1,000 replicates. We used the Pearson product-moment correlation and its 95% confidence interval. In addition, knowing that the correlation is not linear, we measured for each replicate the ratio between the average ancestry of the 50 lowest recombining windows over the average ancestry of the 50 highest recombining windows. We then looked at the distribution of this ratio to obtain its median and 95% confidence interval.

To distinguish between models with the same selection coefficient on average 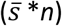 but with different number of variants, we compared for different models the level of genomic spatial correlation of ancestry at the end of the simulations (*i*.*e*. the correlation individual ancestry between two windows a given genetic distance apart). We chose to measure this correlation for different genetic distances (0.01, 0.1, 1 and 10 cM) to see if there is an optimal distance to distinguish between a model with few variants and a high selection coefficient versus a model with more mutation and a low selection coefficient. To do so, we used a custom script that scans chromosome by chromosome and identifies for each 100 kb window a partner window that is at the closest cM distance to the one we defined. We allowed for a difference from the desired distance not greater than half of the target distance. Once all the pairs of windows were identified, we measured for each of a model’s replicates the correlation between all the pairs using a Pearson correlation, which gave us a distribution of correlation coefficients for each scenario.

## Acknowledgments

We thank the University of Wisconsin-Madison Center for High Throughput Computing (CHTC) for computational resources and assistance. This research was funded by US NIH grant R35 GM136306.

## Notes

### Competing Interest Statement

The authors have declared no competing interest.

