## Supplementary_material for "Interactions between natural selection and recombination shape the genomic landscape of introgression"

September 2021

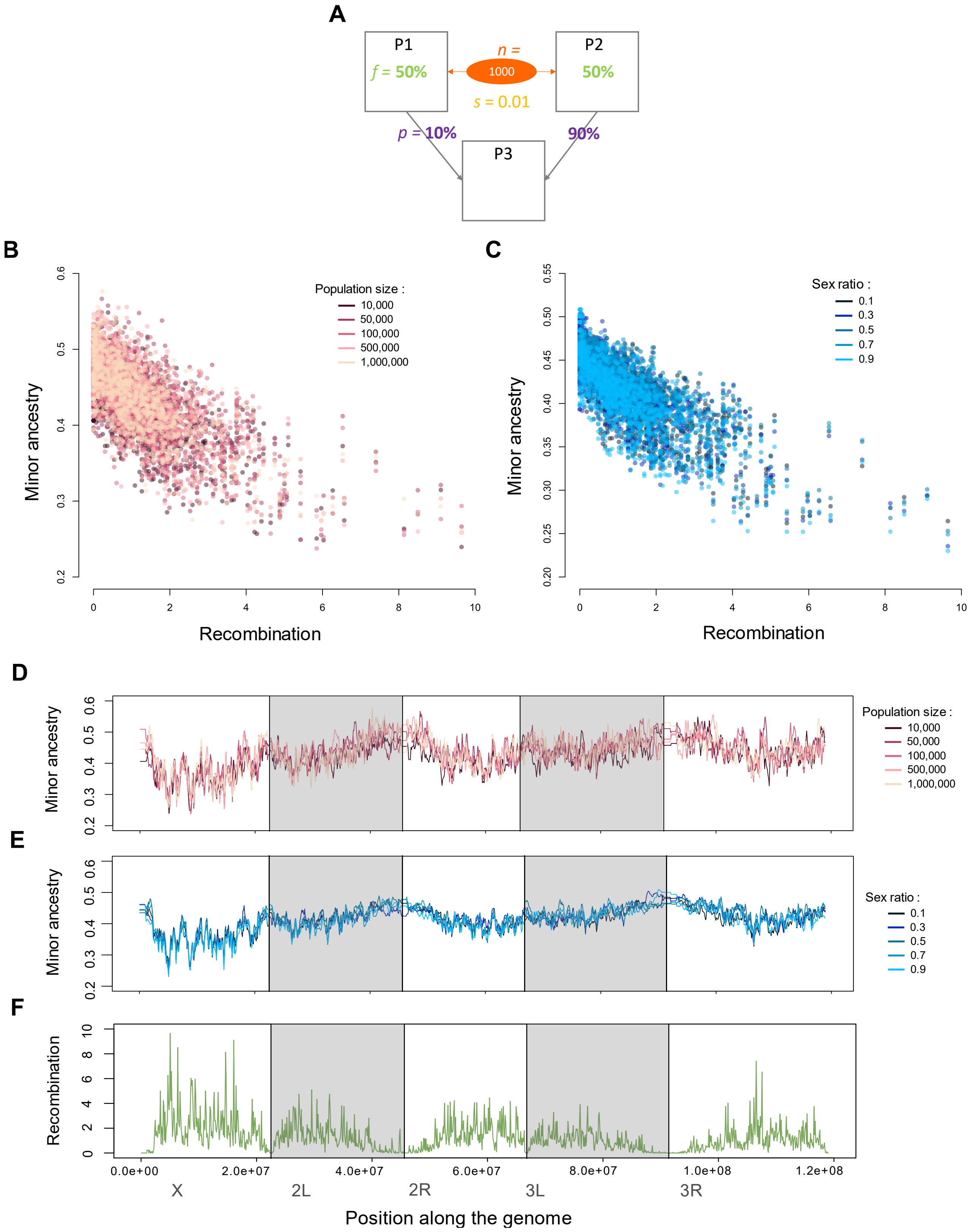


Supplementary figure 1 : **Impact of population sizes and sex-ratio on the relationship between minor ancestry and recombination in a model of directional selection. A.** Schematic representation of the simulated model. Populations p1 and p2 each contain 500 variants with a selection coefficient following an exponential distribution of mean 0.01, contribute respectively at 10% and 90% to the formation of a third population p3 that evolves on its own for 1,600 generations. **B.** Correlation between minor ancestry and recombination for different population size. Each model was replicated 200 times. **C.** Correlation between minor ancestry and recombination for different sex-ratio. Each model was replicated 1,000 times. **D.** Variation in minor ancestry along the genome for different population size. Grey boxes represent the different chromosomes and chromosomes arms **E.** Variation in minor ancestry along the genome for different sex-ratio. **F.** Variation in recombination rate along the genome.

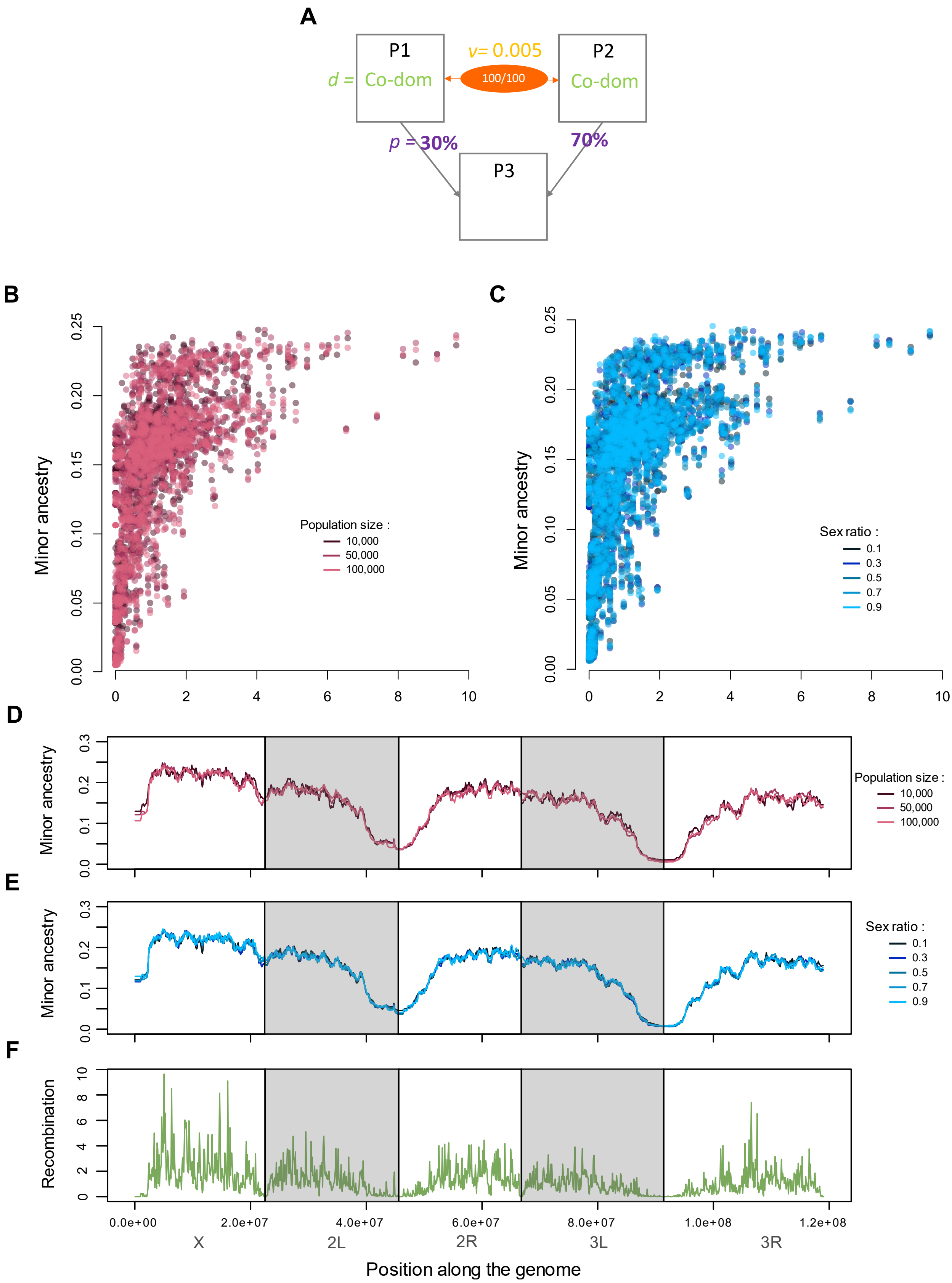


Supplementary figure 2 : **Impact of population sizes and sex-ratio on the relationship between minor ancestry and recombination in a model of pairwise incompatibility. A.** Schematic representation of the simulated model. Populations P1 contains 100 neutral variants involved in a co-dominant pairwise negative interaction with one of 100 alleles fixed within the P2 population. Each negative interaction reduces the fitness of the individual who carries it by a certain value drawn from an exponential distribution with a predetermined mean value of 0.005. P1 and P2 contribute respectively at 30% and 70% to the formation of a third population P3 that evolves on its own for 1,600 generations. **B.** Correlation between minor ancestry and recombination for different population size. Each model was replicated 200 times. **C.** Correlation between minor ancestry and recombination for different sex-ratio. Each model was replicated 1,000 times. **D.** Variation in minor ancestry along the genome for different population size. Grey boxes represent the different chromosomes and chromosomes arms **E.** Variation in minor ancestry along the genome for different sex-ratio. **F.** Variation in recombination rate along the genome.

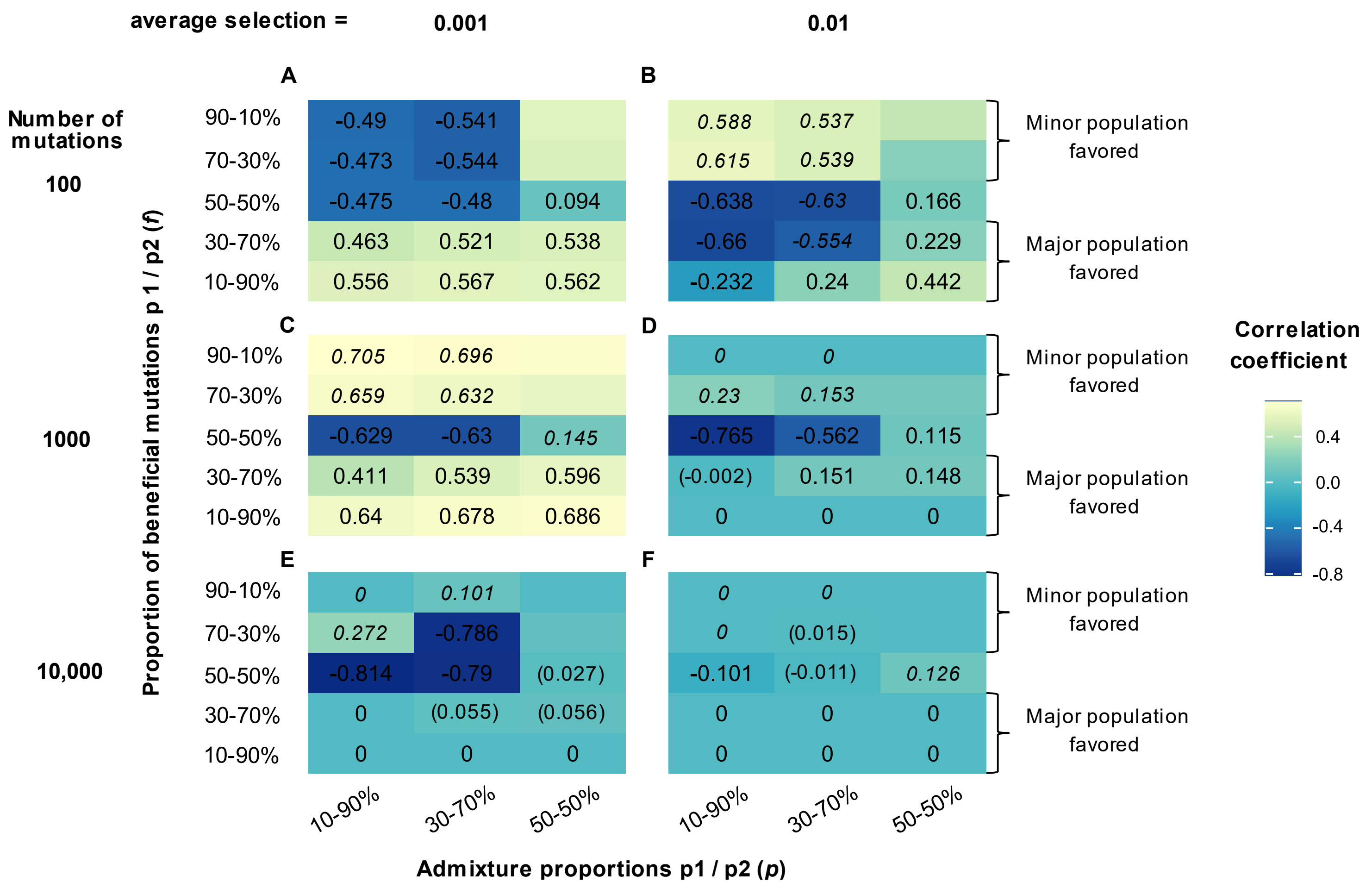


Supplementary Figure 3 : **Correlation coefficient between minor ancestry and recombination for all the model of directional selection measured on the autosomes.** For each grid the value on the left indicates the proportion of beneficial variant present within population p1 and p2 respectively and the value at the bottom the admixture proportion for p1 and p2 respectively. The color indicates if the relationship is negative (darker color) or positive (lighter color). If p2 is the minor ancestry the correlation coefficient is written in italics and if it is non-significant it is written in parentheses. For cases where admixture proportions are 50%-50% we did not simulated cases where the proportion of beneficial variants are 70%-30% and 90%-10%. Indeed, results are expected to be the same as cases where the proportion of beneficial variants are 30%-70% and 10%-90% except with the opposite population as minor ancestry this is why we used the same color for each box but don't have a correlation coefficient. The difference between each grid is the number of beneficial variants present in the simulations and their average selection coefficient. **A.** 100 variants with an average selection coefficient of 0.001, **B.** 100 variants with an average selection coefficient of 0.01, **C.** 1,000 variants with an average selection coefficient of 0.001, **D.** 1,000 variants with an average selection coefficient of 0.01, **E.** 10,000 variants with an average selection coefficient of 0.001 and **F.** 10,000 variants with an average selection coefficient of 0.01.

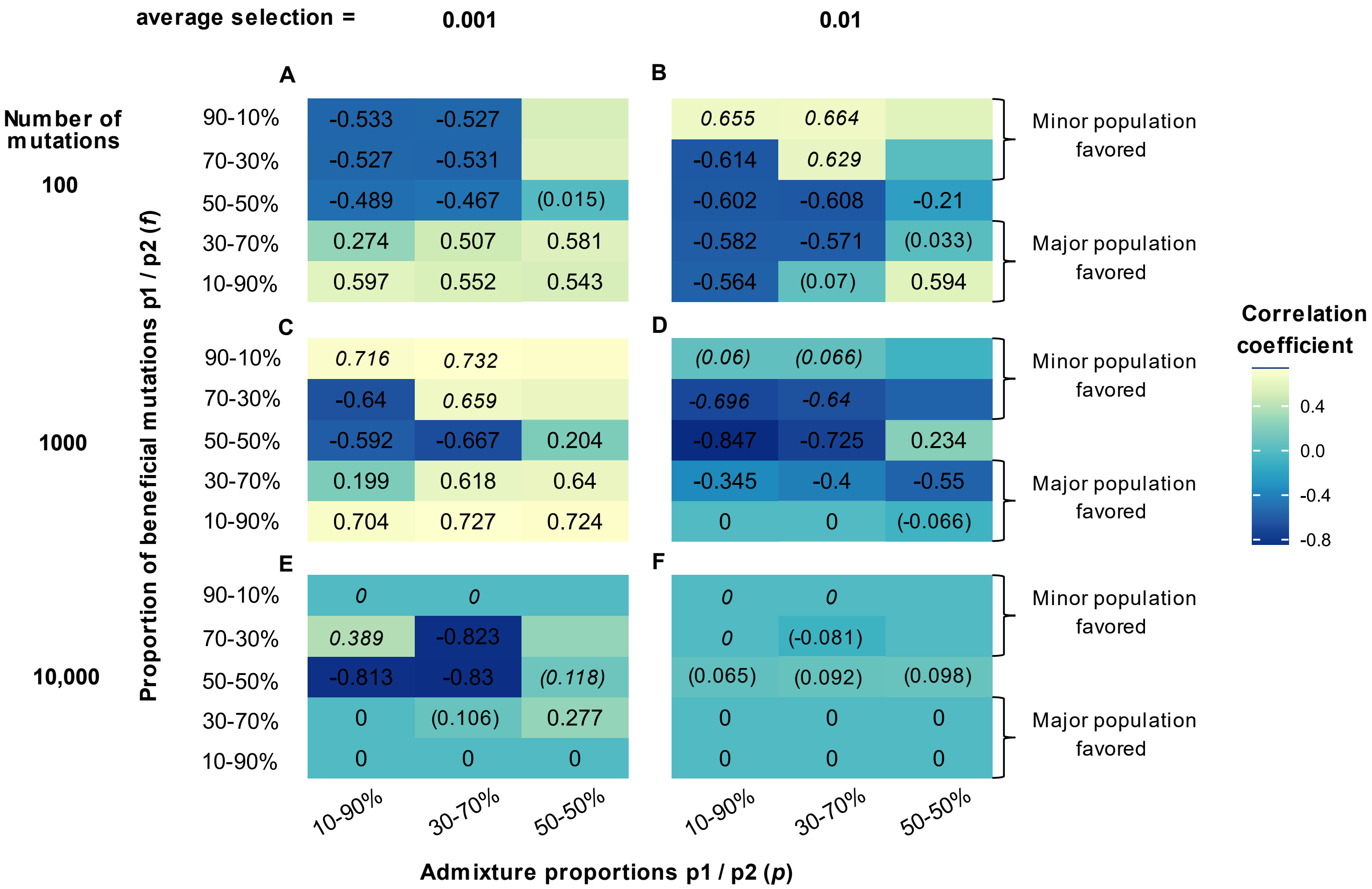


Supplementary Figure 4 : **Correlation coefficient between minor ancestry and recombination for all the model of directional selection measured on the X chromosome.** For each grid the value on the left indicates the proportion of beneficial variant present within population p1 and p2 respectively and the value at the bottom the admixture proportion for p1 and p2 respectively. The color indicates if the relationship is negative (darker color) or positive (lighter color). If p2 is the minor ancestry the correlation coefficient is written in italics and if it is non-significant it is written in parentheses. For cases where admixture proportions are 50%-50% we did not simulated cases where the proportion of beneficial variants are 70%-30% and 90%-10%. Indeed, results are expected to be the same as cases where the proportion of beneficial variants are 30%-70% and 10%-90% except with the opposite population as minor ancestry this is why we used the same color for each box but don't have a correlation coefficient. The difference between each grid is the number of beneficial variants present in the simulations and their average selection coefficient. **A.** 100 variants with an average selection coefficient of 0.001, **B.** 100 variants with an average selection coefficient of 0.01, **C.** 1,000 variants with an average selection coefficient of 0.001, **D.** 1,000 variants with an average selection coefficient of 0.01, **E.** 10,000 variants with an average selection coefficient of 0.001 and **F.** 10,000 variants with an average selection coefficient of 0.01.

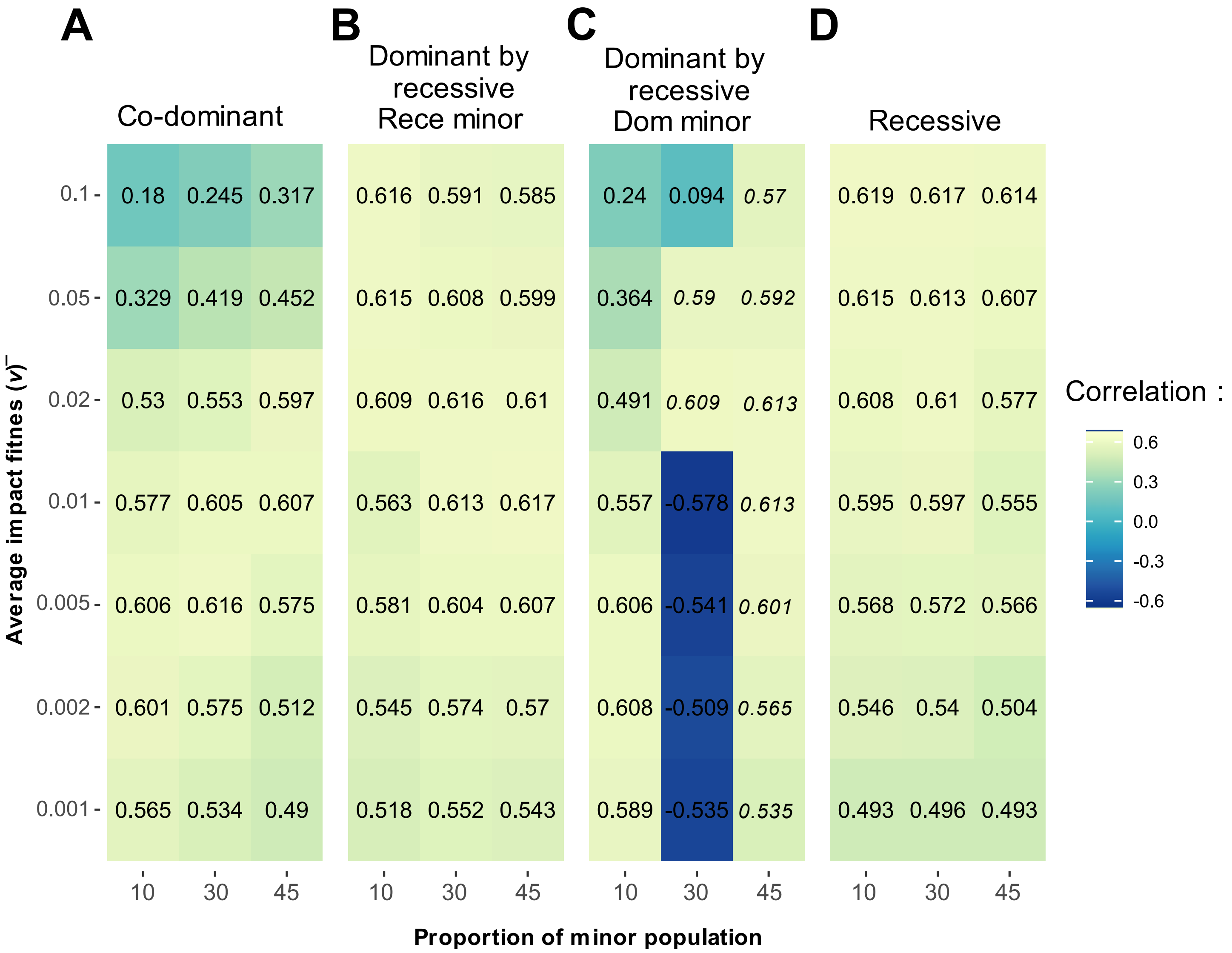


Supplementary Figure 5 : **Correlation coefficient between minor ancestry and recombination for all the model of pairwise incompatibilities measured on the autosomes.** For each grid the value on the left indicates the average fitness impact of each incompatibility ($\bar{v}$) and the value at the bottom the proportion of the minor population at the beginning of the simulations. The color indicates if the relationship is negative (darker color) or positive (lighter color). If the minor ancestry is different at the beginning and the end of the simulations the correlation coefficient is written in italics and if it is non-significant it is written in brackets. The difference between each grid correspond to the dominance of the variants **A.** co-dominant, **B.** dominant by recessive the major population carrying the dominant variants, **C.** dominant by recessive the major population carrying the recessive variants and **D.** recessive.

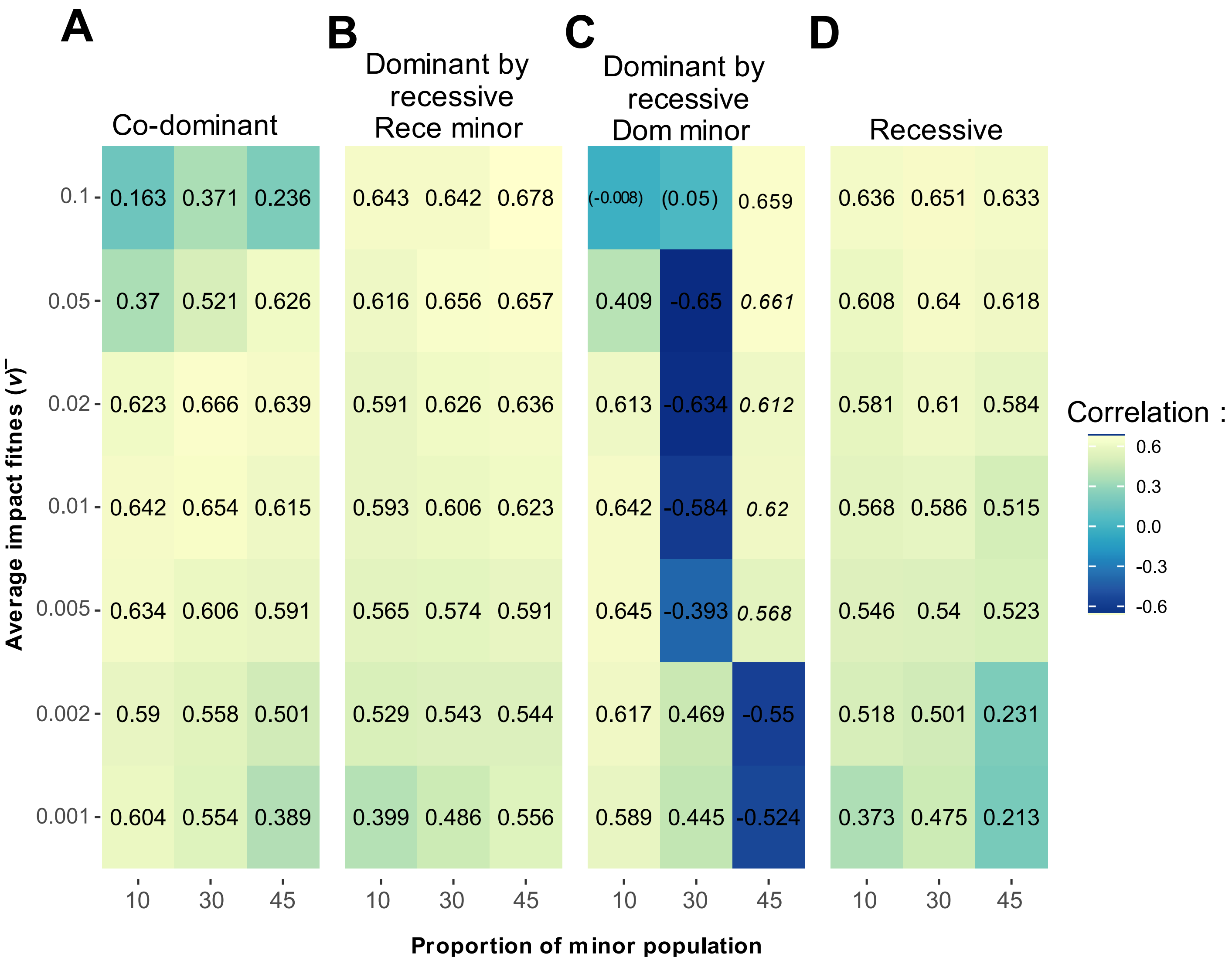


Supplementary Figure 6 : **Correlation coefficient between minor ancestry and recombination for all the model of pairwise incompatibilities measured on the X chromosome.** For each grid the value on the left indicates the average fitness impact of each incompatibility ($\bar{v}$) and the value at the bottom the proportion of the minor population at the beginning of the simulations. The color indicates if the relationship is negative (darker color) or positive (lighter color). If the minor ancestry is different at the beginning and the end of the simulations the correlation coefficient is written in italics and if it is non-significant it is written in brackets. The difference between each grid correspond to the dominance of the variants **A.** co-dominant, **B.** dominant by recessive the major population carrying the dominant variants, **C.** dominant by recessive the major population carrying the recessive variants and **D.** recessive.

| Number of variants (*n*) | Mean of the exponentiel distribution of selection coefficients ($\bar{s}$) | Admixture proportions (*p*) | Fraction of beneficial variants (*f*) p1/p2 | Population with the higher fitness | Minor ancestry after 1,600 generations and proportions | Ratio between least and most recombining windows | Correlation between average minor ancestry and recombination |
|---|---|---|---|---|---|---|---|
| 1000 | 0.001 | 10%- 90% | 10%- 90% | p2 | x : p1 - 1<br>2-3 : p1 -1 | x : 0.20 ± 0.051<br>2-3 : 0.25 ± 0.050 | x : 0.704 $IC_{95\%}$[0.63 ; 0.76] ***<br>2-3 : 0.640 $IC_{95\%}$[0.63 ; 0.76] *** |
| | | | 30%- 70% | p2 | x : p1 -1<br>2-3 : p1 -1 | x : 0.51 ± 0.098<br>2-3 : 0.24 ± 0.062 | x : 0.199 $IC_{95\%}$[0.07 ; 0.32] *<br>2-3 : 0.411 $IC_{95\%}$[0.36 ; 0.46] *** |
| | | | 50%- 50% | None | x : p1 -1<br>2-3 : p1 - 1 | x : 1.61 ± 0.079<br>2-3 : 2.42 ± 0.088 | x : -0.592 $IC_{95\%}$[-0.67 ; -0.50] ***<br>2-3 : -0.629 $IC_{95\%}$[-0.66 ; -0.59] *** |
| | | | 70%- 30% | p1 | x : p1 - 0.985<br>2-3 : p2 - 0.997 | x : 2.00 ± 0.031<br>2-3 : 0.22 ± 0.011 | x : -0.640 $IC_{95\%}$[-0.71 ; -0.55] ***<br>2-3 : 0.659 $IC_{95\%}$[0.62 ; 0.69] *** |
| | | | 90%- 10% | p1 | x : p2 - 1<br>2-3 : p2 - 1 | x : 0.28 ± 0.006<br>2-3 : 0.07 ± 0.004 | x : 0.716 $IC_{95\%}$[0.65 ; 0.77] ***<br>2-3 : 0.705 $IC_{95\%}$[0.67 ; 0.73] *** |
| | | 30%- 70% | 10%- 90% | p2 | x : p1 - 1<br>2-3 : p1 -1 | x : 0.23 ± 0.016<br>2-3 : 0.08 ± 0.015 | x : 0.727 $IC_{95\%}$[0.66 ; 0.78] ***<br>2-3 : 0.678 $IC_{95\%}$[0.64 ; 0.71] *** |
| | | | 30%- 70% | p2 | x : p1 -1<br>2-3 : p1 -1 | x : 0.53 ± 0.042<br>2-3 : 0.24 ± 0.033 | x : 0.618 $IC_{95\%}$[0.53 ; 0.69] ***<br>2-3 : 0.539 $IC_{95\%}$[0.49 ; 0.58] *** |
| | | | 50%- 50% | None | x : p1 -1<br>2-3 : p1 – 0.995 | x : 1.15 ± 0.030<br>2-3 : 1.30 ± 0.035 | x : -0.667 $IC_{95\%}$[-0.73 ; -0.59] ***<br>2-3 : -0.630 $IC_{95\%}$[-0.67 ; -0.59] *** |
| | | | 70%- 30% | p1 | x : p2 -1<br>2-3 : p2 -1 | x : 0.51± 0.017<br>2-3 : 0.22 ± 0.013 | x : 0.659 $IC_{95\%}$[0.58 ; 0.73] ***<br>2-3 : 0.632 $IC_{95\%}$[0.59 ; 0.67] *** |
| | | | 90%- 10% | p1 | x : p2 -1<br>2-3 : p2 -1 | x : 0.26 ± 0.010<br>2-3 : 0.06 ± 0.007 | x : 0.732 $IC_{95\%}$[0.66 ; 0.79] ***<br>2-3 : 0.696 $IC_{95\%}$[0.66 ; 0.73] *** |
| | | 50%- 50% | 10%- 90% | p2 | x : p1 -1<br>2-3 : p1 -1 | x : 0.25 ± 0.012 | x : 0.724 $IC_{95\%}$[0.65 ; 0.78] *** |

| | | | | | | | |
|---|---|---|---|---|---|---|---|
| | | | | | | 2-3 : 0.06 ± 0.010 | 2-3 : 0.686 $IC_{95\%}$[0.65 ; 0.72] *** |
| | | | 30%- 70% | p2 | x : p1 -1<br>2-3 : p1 -1 | x : 0.53 ± 0.023<br>2-3 : 0.24 ± 0.021 | x : 0.640 $IC_{95\%}$[0.55 ; 0.71] ***<br>2-3 : 0.596 $IC_{95\%}$[0.55 ; 0.63] *** |
| | | | 50%- 50% | None | x : p1 – 0.529<br>2-3 : p2 – 0.506 | x : 0.79 ± 0.016<br>2-3 : 0.81 ± 0.019 | x : 0.204 $IC_{95\%}$[0.07 ; 0.33] *<br>2-3 : 0.145 $IC_{95\%}$[0.08 ; 0.21] *** |
| | 0.01 | 10%- 90% | 10%- 90% | p2 | x : p1 – 1<br>2-3 : p1– 1 | x : 1.00 ± 0<br>2-3 : 1.00 ± 0 | x : 0<br>2-3 : 0 |
| | | | 30%- 70% | p2 | x : p1 – 1<br>2-3 : p1 – 1 | x : 1.00 ± 0.218<br>2-3 : 1.00 ± 0.069 | x : -0.345 $IC_{95\%}$[-0.45 ; -0.22] ***<br>2-3 : -0.002 $IC_{95\%}$[-0.06 ; 0.06] N |
| | | | 50%- 50% | None | x : p1 – 0.929<br>2-3 : p1 – 0.846 | x : 1.33 ± 0.045<br>2-3 : 1.10 ± 0.039 | x : -0.847 $IC_{95\%}$[-0.88 ; -0.80] ***<br>2-3 : -0.765 $IC_{95\%}$[-0.79 ; -0.74] *** |
| | | | 70%- 30% | p1 | x : p2 – 1<br>2-3 : p2 – 1 | x : 1.00 ± 7573<br>2-3 : 1.00 ± 49.95 | x : -0.696 $IC_{95\%}$[-0.76 ; -0.62] ***<br>2-3 : 0.23 $IC_{95\%}$[0.17 ; 0.29] *** |
| | | | 90%- 10% | p1 | x : p2 – 1<br>2-3 : p2 – 1 | x : 1.00 ± 0<br>2-3 : 1.00 ± 0 | x : 0.060 $IC_{95\%}$[-0.07 ; -0.19] N<br>2-3 : 0 |
| | | 30%- 70% | 10%- 90% | p2 | x : p1 – 1<br>2-3 : p1– 1 | x : 1.00 ± 0<br>2-3 : 1.00 ± 0 | x : 0<br>2-3 : 0 |
| | | | 30%- 70% | p2 | x : p1 – 1<br>2-3 : p1– 1 | x : 1.00 ± 0.546<br>2-3 : 1.00 ± 38.071 | x : -0.400 $IC_{95\%}$[-0.50 ; -0.28] ***<br>2-3 : 0.151 $IC_{95\%}$[0.09 ; 0.21] *** |
| | | | 50%- 50% | None | x : p1 – 0.733<br>2-3 : p1– 0.661 | x : 1.03 ± 0.034<br>2-3 : 0.952 ± 0.033 | x : -0.725 $IC_{95\%}$[-0.78 ; -0.66] ***<br>2-3 : -0.562 $IC_{95\%}$[-0.60 ; -0.52] *** |
| | | | 70%- 30% | p1 | x : p2 – 1<br>2-3 : p2– 1 | x : 1.00 ± 252.54<br>2-3 : 1 ± 16.135 | x : -0.64 $IC_{95\%}$[-0.71 ; -0.56] ***<br>2-3 : 0.153 $IC_{95\%}$[0.09 ; 0.21] *** |
| | | | 90%- 10% | p1 | x : p2 – 1<br>2-3 : p2– 1 | x : 1.00 ± 0<br>2-3 : 1.00 ± 0 | x : 0.066 $IC_{95\%}$[-0.06 ; -0.19] N<br>2-3 : 0 |
| | | 50%- 50% | 10%- 90% | p2 | x : p1 – 1<br>2-3 : p1– 1 | x : 1.00 ± 0<br>2-3 : 1.00 ± 0 | x : -0.066 $IC_{95\%}$[-0.19 ; 0.06] N<br>2-3 : 0 |
| | | | 30%- 70% | p2 | x : p1 – 1<br>2-3 : p1– 1 | x : 1.00 ± 10.89<br>2-3 : 1.00 ± 7.074 | x : -0.550 $IC_{95\%}$[-0.63 ; -0.45] ***<br>2-3 : 0.148 $IC_{95\%}$[0.09 ; 0.21] *** |

| | | | | | | | |
|---|---|---|---|---|---|---|---|
| | | | 50%- 50% | None | x : p1 – 0.506<br>2-3 : p1– 0.51 | x : 0.97 ± 0.032<br>2-3 : 0.92 ± 0.030 | x : 0.234 $IC_{95\%}$[0.11 ; 0.35] **<br>2-3 : 0.115 $IC_{95\%}$[0.05 ; 0.18] ** |
| 100 | 0.001 | 10%- 90% | 10%- 90% | p2 | x : p1 – 1<br>2-3 : p1– 1 | x : 0.70 ± 0.029<br>2-3 : 0.44 ± 0.022 | x : 0.597 $IC_{95\%}$[0.50 ; 0.67] ***<br>2-3 : 0.556 $IC_{95\%}$[0.51 ; 0.60] ** |
| | | | 30%- 70% | p2 | x : p1 – 1<br>2-3 : p1– 1 | x : 0.84 ± 0.032<br>2-3 : 0.74 ± 0.038 | x : 0.274 $IC_{95\%}$[0.15 ; 0.39] ***<br>2-3 : 0.463 $IC_{95\%}$[0.41 ; 0.51] *** |
| | | | 50%- 50% | None | x : p1 – 1<br>2-3 : p1– 1 | x : 0.98 ± 0.042<br>2-3 : 1.32 ± 0.059 | x : -0.489 $IC_{95\%}$[-0.58 ; -0.38] ***<br>2-3 : -0.475 $IC_{95\%}$[-0.52 ; -0.42] *** |
| | | | 70%- 30% | p1 | x : p1 – 1<br>2-3 : p1– 1 | x : 1.16 ± 0.042<br>2-3 : 2.10 ± 0.062 | x : -0.527 $IC_{95\%}$[-0.62 ; -0.42] ***<br>2-3 : -0.473 $IC_{95\%}$[-0.52 ; -0.42] *** |
| | | | 90%- 10% | p1 | x : p1 – 1<br>2-3 : p1– 1 | x : 1.41 ± 0.044<br>2-3 : 1.82 ± 0.055 | x : -0.533 $IC_{95\%}$[-0.62 ; -0.43] ***<br>2-3 : -0.49 $IC_{95\%}$[-0.54 ; -0.44] *** |
| | | 30%- 70% | 10%- 90% | p2 | x : p1 – 1<br>2-3 : p1– 1 | x : 0.80 ± 0.016<br>2-3 : 0.49 ± 0.014 | x : 0.552 $IC_{95\%}$[0.45 ; 0.64] ***<br>2-3 : 0.567 $IC_{95\%}$[0.52 ; 0.61] *** |
| | | | 30%- 70% | p2 | x : p1 – 1<br>2-3 : p1– 1 | x : 0.90 ± 0.017<br>2-3 : 0.73 ± 0.021 | x : 0.507 $IC_{95\%}$[0.40 ; 0.60] ***<br>2-3 : 0.521 $IC_{95\%}$[0.47 ; 0.57] *** |
| | | | 50%- 50% | None | x : p1 – 1<br>2-3 : p1– 1 | x : 1.00 ± 0.019<br>2-3 : 1.11 ± 0.024 | x : -0.467 $IC_{95\%}$[-0.56 ; -0.36] ***<br>2-3 : -0.48 $IC_{95\%}$[-0.53 ; -0.36] *** |
| | | | 70%- 30% | p1 | x : p1 – 1<br>2-3 : p1– 1 | x : 1.14 ± 0.017<br>2-3 : 1.41 ± 0.021 | x : -0.531 $IC_{95\%}$[-0.62 ; -0.43] ***<br>2-3 : -0.544 $IC_{95\%}$[-0.59 ; -0.50] *** |
| | | | 90%- 10% | p1 | x : p1 – 1<br>2-3 : p1– 0.915 | x : 1.23 ± 0.017<br>2-3 : 1.61 ± 0.026 | x : -0.527 $IC_{95\%}$[-0.61 ; -0.42] ***<br>2-3 : -0.541 $IC_{95\%}$[-0.58 ; -0.49] *** |
| | | 50%- 50% | 10%- 90% | p2 | x : p1 – 0.999<br>2-3 : p1– 1 | x : 0.83 ± 0.012<br>2-3 : 0.52 ± 0.012 | x : 0.543 $IC_{95\%}$[0.44 ; 0.63] ***<br>2-3 : 0.562 $IC_{95\%}$[0.52 ; 0.60] *** |
| | | | 30%- 70% | p2 | x : p1 – 0.92 | x : 0.90 ± 0.011<br>2-3 : 0.76 ± 0.014 | x : 0.581 $IC_{95\%}$[0.49 ; 0.66] *** |

| | | | | | | | |
|---|---|---|---|---|---|---|---|
| | | | | | 2-3 : p1– 0.98 | | 2-3 : 0.538 $IC_{95\%}$[0.49 ; 0.58] *** |
| | | | 50%- 50% | None | x : p1 – 0.506<br>2-3 : p1– 0.508 | x : 0.90 ± 0.010<br>2-3 : 0.85 ± 0.012 | x : 0.015 $IC_{95\%}$[-0.12 ; 0.15] NS<br>2-3 : 0.094 $IC_{95\%}$[0.03 ; 0.16] * |
| | 0.01 | 10%- 90% | 10%- 90% | p2 | x : p1 – 1<br>2-3 : p1– 1 | x : 0.45 ± 0.480<br>2-3 : 0.40 ± 0.493 | x : -0.564 $IC_{95\%}$[-0.65 ; -0.47] ***<br>2-3 : -0.232 $IC_{95\%}$[-0.29 ; -0.17] *** |
| | | | 30%- 70% | p2 | x : p1 – 1<br>2-3 : p1– 1 | x : 1.16 ± 0.215<br>2-3 : 1.60 ± 0.231 | x : -0.582 $IC_{95\%}$[-0.66 ; -0.49] ***<br>2-3 : -0.660 $IC_{95\%}$[-0.69 ; -0.62] *** |
| | | | 50%- 50% | None | x : p1 – 0.999<br>2-3 : p1– 0.999 | x : 1.57 ± 0.098<br>2-3 : 2.34 ± 0.098 | x : -0.602 $IC_{95\%}$[-0.68 ; 0.51] ***<br>2-3 : -0.638 $IC_{95\%}$[-0.67 ; -0.60] *** |
| | | | 70%- 30% | p1 | x : p1 – 0.966<br>2-3 : p2– 0.605 | x : 1.59 ± 0.042<br>2-3 : 0.69 ± 0.045 | x : -0.614 $IC_{95\%}$[-0.69 ; -0.52] ***<br>2-3 : 0.615 $IC_{95\%}$[0.57 ; 0.65] *** |
| | | | 90%- 10% | p1 | x : p2 – 0.939<br>2-3 : p2– 1 | x : 0.58 ± 0.020<br>2-3 : 0.24 ± 0.018 | x : 0.655 $IC_{95\%}$[0.57 ; 0.72] ***<br>2-3 : 0.588 $IC_{95\%}$[0.54 ; 0.63] *** |
| | | 30%- 70% | 10%- 90% | p2 | x : p1 – 1<br>2-3 : p1– 1 | x : 0.54 ± 0.099<br>2-3 : 0.23 ± 0.151 | x : 0.070 $IC_{95\%}$[-0.06 ; 0.20] NS<br>2-3 : 0.24 $IC_{95\%}$[0.18 ; 0.30] *** |
| | | | 30%- 70% | p2 | x : p1 – 1<br>2-3 : p2– 1 | x : 0.95 ± 0.070<br>2-3 : 1.09 ± 0.090 | x : -0.571 $IC_{95\%}$[-0.65 ; -0.47] ***<br>2-3 : -0.554 $IC_{95\%}$[-0.60 ; -0.51] *** |
| | | | 50%- 50% | None | x : p1 – 0.976<br>2-3 : p2– 0.928 | x : 1.14 ± 0.035<br>2-3 : 1.23 ± 0.041 | x : -0.608 $IC_{95\%}$[-0.68 ; -0.52] ***<br>2-3 : -0.630 $IC_{95\%}$[-0.67 ; -0.59] *** |
| | | | 70%- 30% | p1 | x : p2 – 0.904<br>2-3 : p2– 0.995 | x : 0.75 ± 0.022<br>2-3 : 0.69 ± 0.029 | x : 0.629 $IC_{95\%}$[0.54 ; 0.70] ***<br>2-3 : 0.539 $IC_{95\%}$[0.49 ; 0.58] *** |
| | | | 90%- 10% | p1 | x : p2 – 1<br>2-3 : p2– 1 | x : 0.55 ± 0.032<br>2-3 : 0.25 ± 0.042 | x : 0.664 $IC_{95\%}$[0.58 ; 0.73] ***<br>2-3 : 0.537 $IC_{95\%}$[0.49 ; 0.58] *** |
| | | 50%- 50% | 10%- 90% | p2 | x : p1 – 1<br>2-3 : p1– 1 | x : 0.55 ± 0.032<br>2-3 : 0.25 ± 0.042 | x : 0.594 $IC_{95\%}$[0.50 ; 0.67] ***<br>2-3 : 0.442 $IC_{95\%}$[0.39 ; 0.49] *** |

| | | | | | | | |
|---|---|---|---|---|---|---|---|
| | | | 30%- 70% | p2 | x : p1 – 1<br>2-3 : p1– 1 | x : 0.88 ± 0.039<br>2-3 : 0.83 ± 0.049 | x : 0.033 $IC_{95\%}$[-0.10 ; 0.16] NS<br>2-3 : 0.229 $IC_{95\%}$[0.17 ; 0.29] *** |
| | | | 50%- 50% | None | x : p1 – 0.502<br>2-3 : p1– 0.501 | x : 0.85 ± 0.024<br>2-3 : 0.87 ± 0.028 | x : -0.210 $IC_{95\%}$[-0.33 ; -0.08] **<br>2-3 : 0.166 $IC_{95\%}$[0.10 ; 0.23] *** |
| 10,000 | 0.001 | 10%- 90% | 10%- 90% | p2 | x : p1 – 1<br>2-3 : p1– 1 | x : 1.00 ± 0<br>2-3 : 1.00 ± 0 | x : 0<br>2-3 : 0 |
| | | | 30%- 70% | p2 | x : p1 – 1<br>2-3 : p1– 1 | x : 1.00 ± 0<br>2-3 : 1.00 ± 0 | x : 0<br>2-3 : 0 |
| | | | 50%- 50% | None | x : p1 – 1<br>2-3 : p1– 1 | x : 1 .76 ± 0.059<br>2-3 : 1.67 ± 0.051 | x : -0.813 $IC_{95\%}$[-0.85 ; -0.76] ***<br>2-3 : -0.814 $IC_{95\%}$[-0.83 ; -0.79] *** |
| | | | 70%- 30% | p1 | x : p2 – 1<br>2-3 : p2– 1 | x : 1.00 ± 0.96<br>2-3 : 1.00 ± 0 | x : 0.389 $IC_{95\%}$[0.27 ; 0.49] ***<br>2-3 : 0.272 $IC_{95\%}$[0.21 ; 0.33] *** |
| | | | 90%- 10% | p1 | x : p2 – 1<br>2-3 : p2– 1 | x : 1.00 ± 0<br>2-3 : 1.00 ± 0 | x : 0<br>2-3 : 0 |
| | | 30%- 70% | 10%- 90% | p2 | x : p1 – 1<br>2-3 : p1– 1 | x : 1.00 ± 0<br>2-3 : 1.00 ± 0 | x : 0<br>2-3 : 0 |
| | | | 30%- 70% | p2 | x : p1 – 1<br>2-3 : p1– 1 | x : 1.00 ± 0.006<br>2-3 : 1.00 ± 0 | x : 0.106 $IC_{95\%}$[-0.02 ; 0.23] NS<br>2-3 : 0.055 $IC_{95\%}$[-0.01 ; 0.12] NS |
| | | | 50%- 50% | None | x : p1 – 0.911<br>2-3 : p1– 0.864 | x : 1 .12 ± 0.030<br>2-3 : 1.06 ± 0.30 | x : -0.830 $IC_{95\%}$[-0.87 ; -0.78] ***<br>2-3 : -0.79 $IC_{95\%}$[-0.81 ; -0.76] *** |
| | | | 70%- 30% | p1 | x : p1 – 0.922<br>2-3 : p1– 0.849 | x : 1 .12 ± 0.030<br>2-3 : 1.08 ± 0.032 | x : -0.823 $IC_{95\%}$[-0.86 ; -0.77] ***<br>2-3 : -0.786 $IC_{95\%}$[-0.81 ; -0.76] *** |
| | | | 90%- 10% | p1 | x : p2 – 1<br>2-3 : p2– 1 | x : 1.00 ± 0<br>2-3 : 1.00 ± 0 | x : 0<br>2-3 : 0.101 $IC_{95\%}$[0.04 ; 0.16] * |
| | | 50%- 50% | 10%- 90% | p2 | x : p1 – 1<br>2-3 : p1– 1 | x : 1.00 ± 0<br>2-3 : 1.00 ± 0 | x : 0<br>2-3 : 0 |
| | | | 30%- 70% | p2 | x : p1 – 1<br>2-3 : p1– 1 | x : 1 .00 ± 0.171<br>2-3 : 1.00 ± 0 | x : 0.277 $IC_{95\%}$[0.15 ; 0.39] ***<br>2-3 : 0.056 $IC_{95\%}$[-0.01 ; 0.12] NS |
| | | | 50%- 50% | None | x : p2 – 0.523<br>2-3 : p1– 0.516 | x : 0.86 ± 0.022<br>2-3 : 0.88 ± 0.023 | x : 0.118 $IC_{95\%}$[-0.01 ; 0.25] NS<br>2-3 : 0.027 $IC_{95\%}$[-0.03 ; 0.09] NS |
| | 0.01 | 10%- 90% | 10%- 90% | p2 | x : p1 – 1<br>2-3 : p1– 1 | x : 1.00 ± 0<br>2-3 : 1.00 ± 0 | x : 0<br>2-3 : 0 |

| | | | | | | | |
|---|---|---|---|---|---|---|---|
| | | | 30%- 70% | p2 | x : p1 – 1<br>2-3 : p1– 1 | x : 1.00 ± 0<br>2-3 : 1.00 ± 0 | x : 0<br>2-3 : 0 |
| | | | 50%- 50% | None | x : p1 – 0.655<br>2-3 : p1– 0.594 | x : 0.87 ± 0.026<br>2-3 : 0.83 ± 0.029 | x : 0.065 IC$_{95\%}$[-0.07 ; 0.19] NS<br>2-3 : -0.101 IC$_{95\%}$[-0.16 ; -0.04] * |
| | | | 70%- 30% | p1 | x : p2 – 1<br>2-3 : p2– 1 | x : 1.00 ± 0<br>2-3 : 1.00 ± 0 | x : 0<br>2-3 : 0 |
| | | | 90%- 10% | p1 | x : p2 – 1<br>2-3 : p2– 1 | x : 1.00 ± 0<br>2-3 : 1.00 ± 0 | x : 0<br>2-3 : 0 |
| | | 30%- 70% | 10%- 90% | p2 | x : p1 – 1<br>2-3 : p1– 1 | x : 1.00 ± 0<br>2-3 : 1.00 ± 0 | x : 0<br>2-3 : 0 |
| | | | 30%- 70% | p2 | x : p1 – 1<br>2-3 : p1– 1 | x : 1.00 ± 0<br>2-3 : 1.00 ± 0 | x : 0<br>2-3 : 0 |
| | | | 50%- 50% | None | x : p1 – 0.572<br>2-3 : p1– 0.546 | x : 0.832 ± 0.024<br>2-3 : 0.842 ± 0.026 | x : 0.092 IC$_{95\%}$[-0.04 ; 0.22] NS<br>2-3 : -0.011 IC$_{95\%}$[-0.07 ; 0.05] NS |
| | | | 70%- 30% | p1 | x : p1 – 0.564<br>2-3 : p1– 0.536 | x : 0.84 ± 0.024<br>2-3 : 0.83 ± 0.028 | x : -0.081 IC$_{95\%}$[-0.21 ; 0.05] NS<br>2-3 : 0.015 IC$_{95\%}$[-0.05 ; 0.08] NS |
| | | | 90%- 10% | p1 | x : p2 – 1<br>2-3 : p2– 1 | x : 1 00 ± 0<br>2-3 : 1.00 ± 0 | x : 0<br>2-3 : 0 |
| | | 50%- 50% | 10%- 90% | p2 | x : p1 – 1<br>2-3 : p1– 1 | x : 1.00 ± 0<br>2-3 : 1.00 ± 0 | x : 0<br>2-3 : 0 |
| | | | 30%- 70% | p2 | x : p1 – 1<br>2-3 : p1– 1 | x : 1.00 ± 0<br>2-3 : 1.00 ± 0 | x : 0<br>2-3 : 0 |
| | | | 50%- 50% | None | x : p1 – 0.516<br>2-3 : p2– 0.503 | x : 0.85 ± 0.022<br>2-3 : 0.81 ± 0.026 | x : 0.098 IC$_{95\%}$[-0.03 ; 0.23] NS<br>2-3 : 0.126 IC$_{95\%}$[0.06 ; 0.19] *** |

Supplementary Table 1 : **Results of all the simulations under the model of directional selection.** Each model is characterized by 4 parameters, *n* the number of positively selected variants present in the simulation which selection coefficient follows an exponential distribution of average $\bar{S}$ and are distributed between the two populations p1 and p2 in proportion $f$ . P1 and p2 are mixed in proportion *p* to create a third population. For each set of parameter, we indicate the fitter population, the minority ancestry after 1,600 generations on the sex chromosome and the autosomes as well as the proportions of model (over 1,000) in which this ancestry is minority. The median of the distribution of the ratio between the 50 lowest and highest recombining regions for all replicates as well as the 95% confidence interval of the value separately for the sex chromosome and the autosomes. Finally, the correlation coefficient between the average (across the 1,000 replicates) minor ancestry and the recombination rates for the X chromosome and the autosomes. The significance of the correlation is shown with different symbols, NS if the p-value is >0.05, * if 0.05>p-value>0.001, **if 0.001>p-value>2.2e-16 and *** if p-value < 2.2e-16.

| Average selection genome wide | Admixture proportions (*p*) | Fraction of beneficial variants (*f*) p1/p2 | Average distance between windows (cM) | Spatial correlation coefficient of p2 ancestry between windows |
|---|---|---|---|---|
| 1 | 10%- 90% | 10%- 90% | 0.0115 | Few: 0.952 ± 0.008<br>Many: 0.967 ± 0.009 |
| | | | 0.114 | Few: 0.694 ± 0.010<br>Many: 0.475 ± 0.009 |
| | | | 1.016 | Few: 0.077 ± 0.006<br>Many: 0.062 ± 0.004 |
| | | | 10.018 | Few: 0.001 ± 0.003<br>Many: 0.019 ± 0.003 |
| | | 30% - 70% | 0.0115 | Few: 0.948 ± 0.006<br>Many: 0.906 ± 0.010 |
| | | | 0.114 | Few: 0.757 ± 0.006<br>Many: 0.601 ± 0.007 |
| | | | 1.016 | Few: 0.165 ± 0.007<br>Many: 0.078 ± 0.004 |
| | | | 10.018 | Few: -0.003 ± 0.004<br>Many: 0.008 ± 0.003 |
| | | 50% - 50% | 0.0115 | Few: 0.952 ± 0.003<br>Many: 0.961 ± 0.002 |
| | | | 0.114 | Few: 0.822 ± 0.004<br>Many: 0.719 ± 0.004 |
| | | | 1.016 | Few: 0.339 ± 0.008<br>Many: 0.164 ± 0.007 |
| | | | 10.018 | Few: 0.021 ± 0.007<br>Many: 0.001 ± 0.004 |
| | | 70% - 30% | 0.0115 | Few: 0.950 ± 0.004<br>Many: 0.965 ± 0.002 |
| | | | 0.114 | Few: 0.853 ± 0.003<br>Many: 0.820 ± 0.002 |
| | | | 1.016 | Few: 0.483 ± 0.006<br>Many: 0.442 ± 0.005 |
| | | | 10.018 | Few: 0.088 ± 0.008<br>Many: 0.091 ± 0.006 |
| | | 90% - 10% | 0.0115 | Few: 0.962 ± 0.004<br>Many: 0.927 ± 0.006 |
| | | | 0.114 | Few: 0.849 ± 0.003<br>Many: 0.796 ± 0.002 |
| | | | 1.016 | Few: 0.545 ± 0.005<br>Many: 0.513 ± 0.003 |
| | | | 10.018 | Few: 0.166 ± 0.008<br>Many: 0.274 ± 0.005 |
| | 30% - 70% | 10% - 90% | 0.0115 | Few: 0.938 ± 0.008<br>Many: 0.898 ± 0.013 |
| | | | 0.114 | Few: 0.711 ± 0.008<br>Many: 0.544 ± 0.006 |
| | | | 1.016 | Few: 0.144 ± 0.006<br>Many: 0.141 ± 0.004 |

| | | | | |
|---|---|---|---|---|
| | | | 10.018 | Few: 0.010 ± 0.004<br>Many: 0.059 ± 0.004 |
| | | 30% - 70% | 0.0115 | Few: 0.948 ± 0.004<br>Many: 0.933 ± 0.006 |
| | | | 0.114 | Few: 0.779 ± 0.005<br>Many: 0.664 ± 0.005 |
| | | | 1.016 | Few: 0.263 ± 0.007<br>Many: 0.156 ± 0.004 |
| | | | 10.018 | Few: 0.019 ± 0.006<br>Many: 0.031 ± 0.004 |
| | | 50% - 50% | 0.0115 | Few: 0.947 ± 0.003<br>Many: 0.960 ± 0.002 |
| | | | 0.114 | Few: 0.820 ± 0.003<br>Many: 0.728 ± 0.004 |
| | | | 1.016 | Few: 0.392 ± 0.007<br>Many: 191 ± 0.006 |
| | | | 10.018 | Few: 0.044 ± 0.008<br>Many: 0.008 ± 0.005 |
| | | 70% - 30% | 0.0115 | Few: 0.049 ± 0.004<br>Many: 0.956 ± 0.003 |
| | | | 0.114 | Few: 0.823 ± 0.004<br>Many: 0.763 ± 0.003 |
| | | | 1.016 | Few: 0.418 ± 0.006<br>Many: 0.334 ± 0.005 |
| | | | 10.018 | Few: 0.075 ± 0.008<br>Many: 0.079 ± 0.005 |
| | | 90% - 10% | 0.0115 | Few: 0.947 ± 0.005<br>Many: 0.879 ± 0.011 |
| | | | 0.114 | Few: 0.770 ± 0.006<br>Many: 0.687 ± 0.003 |
| | | | 1.016 | Few: 0.359 ± 0.007<br>Many: 0.347 ± 0.004 |
| | | | 10.018 | Few: 0.097 ± 0.006<br>Many: 0.179 ± 0.005 |
| | 50% - 50% | 10% - 90% | 0.0115 | Few: 0.944 ± 0.006<br>Many: 0.864 ± 0.013 |
| | | | 0.114 | Few: 0.734 ± 0.007<br>Many: 0.607 ± 0.005 |
| | | | 1.016 | Few: 0.238 ± 0.007<br>Many: 0.225 ± 0.004 |
| | | | 10.018 | Few: 0.045 ± 0.005<br>Many: 0.108 ± 0.004 |
| | | 30% - 70% | 0.0115 | Few: 0945 ± 0.004<br>Many: 0.947 ± 0.004 |
| | | | 0.114 | Few: 0.799 ± 0.004<br>Many: 0.712 ± 0.004 |
| | | | 1.016 | Few: 0.343 ± 0.007<br>Many: 0.241 ± 0.005 |
| | | | 10.018 | Few: 0.044 ± 0.007<br>Many: 0.060 ± 0.005 |
| | | 50% - 50% | 0.0115 | Few: 0.946 ± 0.004 |

| | | | | |
|---|---|---|---|---|
| | | | | Many: 0.958 ± 0.002 |
| | | | 0.114 | Few: 0.821 ± 0.003<br>Many: 0.727 ± 0.003 |
| | | | 1.016 | Few: 0.402 ± 0.006<br>Many: 0.192 ± 0.005 |
| | | | 10.018 | Few: 0.055 ± 0.008<br>Many: 0.013 ± 0.005 |
| 10 | 10% - 90% | 10% - 90% | 0.0115 | Few: 1<br>Many: 1 |
| | | | 0.114 | Few: 1<br>Many: 1 |
| | | | 1.016 | Few: 1<br>Many: 1 |
| | | | 10.018 | Few: 1.009 ± 0.018<br>Many: 1.010 ± 0.017 |
| | | 30% - 70% | 0.0115 | Few: 0.999 ± 0.0002<br>Many: 1 |
| | | | 0.114 | Few: 0.975 ± 0.008<br>Many: 1 |
| | | | 1.016 | Few: 0.895 ± 0.018<br>Many: 1 |
| | | | 10.018 | Few: 0.936 ± 0.024<br>Many: 1.009 ± 0.017 |
| | | 50% - 50% | 0.0115 | Few: 0.840 ± 0.005<br>Many: 0.637 ± 0.003 |
| | | | 0.114 | Few: 0.643 ± 0.004<br>Many: 0.013 ± 0.005 |
| | | | 1.016 | Few: 0.196 ± 0.004<br>Many: 0.167 ± 0.004 |
| | | | 10.018 | Few: 0.027 ± 0.020<br>Many: 0.027 ± 0.020 |
| | | 70% - 30% | 0.0115 | Few: 0.990 ± 0.003<br>Many: 1 |
| | | | 0.114 | Few: 0.731 ± 0.012<br>Many: 0.798 ± 0.023 |
| | | | 1.016 | Few: 0.156 ± 0.011<br>Many: 0.126 ± 0.020 |
| | | | 10.018 | Few: 0.048 ± 0.023<br>Many: 0.229 ± 0.032 |
| | | 90% - 10% | 0.0115 | Few: 1<br>Many: 1 |
| | | | 0.114 | Few: 1<br>Many: 1 |
| | | | 1.016 | Few: 0.999 ± 0.002<br>Many: 1 |
| | | | 10.018 | Few: 1.008. ± 0.018<br>Many: 1.009 ± 0.018 |
| | 30% - 70% | 10% - 90% | 0.0115 | Few: 1<br>Many: 1 |
| | | | 0.114 | Few: 1<br>Many: 1 |

| | | | | |
|---|---|---|---|---|
| | | | 1.016 | Few: 1<br>Many: 1 |
| | | | 10.018 | Few: 1<br>Many: 1 |
| | | 30% - 70% | 0.0115 | Few: 0.999 ± 0.001<br>Many: 1 |
| | | | 0.114 | Few: 0.941 ± 0.010<br>Many: 0.992 ± 0.005 |
| | | | 1.016 | Few: 0.659 ± 0.028<br>Many: 0.913 ± 0.017 |
| | | | 10.018 | Few: 0.730 ± 0.028<br>Many: 0.935 ± 0.015 |
| | | 50% - 50% | 0.0115 | Few: 0.831 ± 0.006<br>Many: 0.880 ± 0.004 |
| | | | 0.114 | Few: 0.624 ± 0.003<br>Many: 0.605 ± 0.003 |
| | | | 1.016 | Few: 0.187 ± 0.004<br>Many: 0.146 ± 0.004 |
| | | | 10.018 | Few: 0.013 ± 0.004<br>Many: 0.010 ± 0.004 |
| | | 70% - 30% | 0.0115 | Few: 0.997 ± 0.002<br>Many: 0.873 ± 0.005 |
| | | | 0.114 | Few: 0.808 ± 0.015<br>Many: 0.608 ± 0.003 |
| | | | 1.016 | Few: 0.269 ± 0.023<br>Many: 0.149 ± 0.004 |
| | | | 10.018 | Few: 0.283 ± 0.028<br>Many: 0.010 ± 0.004 |
| | | 90% - 10% | 0.0115 | Few: 1<br>Many: 1 |
| | | | 0.114 | Few: 1<br>Many: 1 |
| | | | 1.016 | Few: 0.999 ± 0.002<br>Many: 0.999 ± 0.002 |
| | | | 10.018 | Few: 1<br>Many: 1 |
| | 50% - 50% | 10% - 90% | 0.0115 | Few: 1<br>Many: 1 |
| | | | 0.114 | Few: 1<br>Many: 1 |
| | | | 1.016 | Few: 0.999 ± 0.002<br>Many: 1 |
| | | | 10.018 | Few: 0.999 ± 0.002<br>Many: 1 |
| | | 30% - 70% | 0.0115 | Few: 0.998 ± 0.002<br>Many: 1 |
| | | | 0.114 | Few: 0.879 ± 0.135<br>Many: 0.983 ± 0.007 |
| | | | 1.016 | Few: 0.446 ± 0.028<br>Many: 0.797± 0.025 |
| | | | 10.018 | Few: 0.858 ± 0.022 |

| | | | | |
|---|---|---|---|---|
| | | | | Many: 0.999 ± 0.002 |
| | | 50% - 50% | 0.0115 | Few: 0.826 ± 0.006<br>Many: 0.870± 0.005 |
| | | | 0.114 | Few: 0.623 ± 0.003<br>Many: 0.601 ± 0.003 |
| | | | 1.016 | Few: 0.183 ± 0.004<br>Many: 0.150 ± 0.004 |
| | | | 10.018 | Few: 0.012 ± 0.005<br>Many: 0.010 ± 0.004 |

Supplementary Table 2 : **Comparison of spatial correlation between models with the same level of selection genome wide but with either a high number of variants with a low selection coefficient or a low number of variants with a high selection coefficient.** When selection is on average equal to 1, model with many variants have *n*=1,000 variants with an average selection coefficient of $\bar{s}$=0.001 and model with few variants have *n*=100 variants with an average selection coefficient of $\bar{s}$=0.01. When selection is on average equal to 10 model with many variants have *n*=10,000 variants with an average selection coefficient of $\bar{s}$=0.001 and model with few variants have *n*=1,000 variants with an average selection coefficient of $\bar{s}$=0.01. Each model is also characterized by 2 other parameters, the fraction of beneficial variants presents within populations p1 and p2 (*f*) and the proportion in which p1 and p2 are mixed (*p*) to create a third population. For each set of parameters, and four different genomic distances (0.01, 0.1, 1 and 10 cM) we indicate the mean level of spatial correlation of p2 ancestry between all 100kb windows separated by the considered distance and its 95% confidence interval.

| Dominance ($d$) | Admixture proportions ($p$) | Mean of the exponential distribution of fitness impact ($\bar{v}$) | Minor ancestry after 1,600 generations and proportions | Ratio between least and most recombining windows | Correlation between average minor ancestry and recombination |
|---|---|---|---|---|---|
| Co-dominant | 10% - 90% | 0.001 | x : p1 – 1<br>2-3 : p1 -1 | x : 0.75 ± 0.023<br>2-3 : 0.42 ± 0.017 | x : 0.604 $IC_{95\%}$[0.51 ; 0.68] ***<br>2-3 : 0.565 $IC_{95\%}$[0.52 ; 0.61] *** |
| | | 0.002 | x : p1 – 1<br>2-3 : p1 -1 | x : 0.67 ± 0.024<br>2-3 : 0.28 ± 0.015 | x : 0.590 $IC_{95\%}$[0.50 ; 0.67] ***<br>2-3 : 0.601 $IC_{95\%}$[0.56 ; 0.64] *** |
| | | 0.005 | x : p1 – 1<br>2-3 : p1 -1 | x : 0.52 ± 0.027<br>2-3 : 0.18 ± 0.014 | x : 0.634 $IC_{95\%}$[0.55 ; 0.71] ***<br>2-3 : 0.606 $IC_{95\%}$[0.56 ; 0.64] *** |
| | | 0.01 | x : p1 – 1<br>2-3 : p1 -1 | x : 0.46 ± 0.037<br>2-3 : 0.19 ± 0.018 | x : 0.642 $IC_{95\%}$[0.53 ; 0.62] ***<br>2-3 : 0.577 $IC_{95\%}$[0.53 ; 0.62] *** |
| | | 0.02 | x : p1 – 1<br>2-3 : p1 -1 | x : 0.38 ± 0.069<br>2-3 : 0.33 ± 0.051 | x : 0.623 $IC_{95\%}$[0.53 ; 0.70] ***<br>2-3 : 0.530 $IC_{95\%}$[0.48 ; 0.57] *** |
| | | 0.05 | x : p1 – 1<br>2-3 : p1 -1 | x : 0.54 ± 2.466<br>2-3 : 1 ± 7.432 | x : 0.370 $IC_{95\%}$[0.25 ; 0.48] ***<br>2-3 : 0.329 $IC_{95\%}$[0.27 ; 0.38] *** |
| | | 0.1 | x : p1 – 1<br>2-3 : p1 -1 | x : 0.52 ± 0.027<br>2-3 : 0.18 ± 0.014 | x : 0.163 $IC_{95\%}$[0.03 ; 0.29] *<br>2-3 : 0.180 $IC_{95\%}$[0.12 ; 0.24] ** |
| | 30% - 70% | 0.001 | x : p1 – 1<br>2-3 : p1 -1 | x : 0.88 ± 0.014<br>2-3 : 0.62 ± 0.014 | x : 0.554 $IC_{95\%}$[0.46 ; 0.64] ***<br>2-3 : 0.534 $IC_{95\%}$[0.49 ; 0.58] *** |
| | | 0.002 | x : p1 – 1<br>2-3 : p1 -1 | x : 0.78 ± 0.015<br>2-3 : 0.44 ± 0.013 | x : 0.558 $IC_{95\%}$[0.46 ; 0.64] ***<br>2-3 : 0.575 $IC_{95\%}$[0.53 ; 0.62] *** |
| | | 0.005 | x : p1 – 1<br>2-3 : p1 - 1 | x : 0.65 ± 0.019<br>2-3 : 0.23 ± 0.011 | x : 0.606 $IC_{95\%}$[0.51 ; 0.68] ***<br>2-3 : 0.616 $IC_{95\%}$[0.58 ; 0.65] *** |
| | | 0.01 | x : p1 – 1<br>2-3 : p1 - 1 | x : 0.55 ± 0.021<br>2-3 : 0.18 ± 0.010 | x : 0.654 $IC_{95\%}$[0.57 ; 0.72] ***<br>2-3 : 0.605 $IC_{95\%}$[0.56 ; 0.64] *** |
| | | 0.02 | x : p1 – 1<br>2-3 : p1 -1 | x : 0.46 ± 0.029<br>2-3 : 0.18 ± 0.013 | x : 0.666 $IC_{95\%}$[0.59 ; 0.73] ***<br>2-3 : 0.553 $IC_{95\%}$[0.51 ; 0.59] *** |
| | | 0.05 | x : p1 – 1<br>2-3 : p1 -1 | x : 0.34 ± 0.121<br>2-3 : 0.52 ± 0.338 | x : 0.521 $IC_{95\%}$[0.42 ; 0.61] ***<br>2-3 : 0.419 $IC_{95\%}$[0.36 ; 0.47] *** |
| | | 0.1 | x : p1 – 1<br>2-3 : p1 - 1 | x : 1 ± 7.592<br>2-3 : 1 ± 0.362 | x : 0.371 $IC_{95\%}$[0.25 ; 0.48] **<br>2-3 : 0.245 $IC_{95\%}$[0.18 ; 0.30] ** |
| | 45% - 55% | 0.001 | x : p1 – 0.98<br>2-3 : p1 – 0.997 | x : 0.97 ± 0.011<br>2-3 : 0.89 ± 0.014 | x : 0.389 $IC_{95\%}$[0.27 ; 0.46] ***<br>2-3 : 0.490 $IC_{95\%}$[0.44 ; 0.54] *** |
| | | 0.002 | x : p1 – 0.969<br>2-3 : p1 – 0.991 | x : 0.92 ± 0.014<br>2-3 : 0.81 ± 0.019 | x : 0.501 $IC_{95\%}$[0.39 ; 0.59] ***<br>2-3 : 0.512 $IC_{95\%}$[0.46 ; 0.56] *** |
| | | 0.005 | x : p1 – 0.986<br>2-3 : p1 – 0.992 | x : 0.84 ± 0.020<br>2-3 : 0.57 ± 0.024 | x : 0.591 $IC_{95\%}$[0.50 ; 0.67] ***<br>2-3 : 0.575 $IC_{95\%}$[0.53 ; 0.61] *** |
| | | 0.01 | x : p1 – 0.999<br>2-3 : p1 – 0.997 | x : 0.69 ± 0.023<br>2-3 : 0.27 ± 0.023 | x : 0.615 $IC_{95\%}$[0.53 ; 0.69] ***<br>2-3 : 0.607 $IC_{95\%}$[0.56 ; 0.64] *** |
| | | 0.02 | x : p1 – 1<br>2-3 : p1 – 1 | x : 0.51 ± 0.025<br>2-3 : 0.18 ± 0.013 | x : 0.639 $IC_{95\%}$[0.55 ; 0.71] ***<br>2-3 : 0.597 $IC_{95\%}$[0.55 ; 0.64] *** |
| | | 0.05 | x : p1 – 1<br>2-3 : p1 – 1 | x : 0.41 ± 0.054<br>2-3 : 0.28 ± 0.041 | x : 0.626 $IC_{95\%}$[0.54 ; 0.70] ***<br>2-3 : 0.452 $IC_{95\%}$[0.40 ; 0.50] *** |
| | | 0.1 | x : p1 – 1<br>2-3 : p1 - 1 | x : 0.55 ± 3.660<br>2-3 : 1 ± 31.85 | x : 0.236 $IC_{95\%}$[0.11 ; 0.36] **<br>2-3 : 0.317 $IC_{95\%}$[0.26 ; 0.37] *** |
| Recessive | 10% - 90% | 0.001 | x : p1 – 1<br>2-3 : p1 - 1 | x : 0.90 ± 0.024<br>2-3 : 0.81 ± 0.019 | x : 0.373 $IC_{95\%}$[0.25 ; 0.48] **<br>2-3 : 0.493 $IC_{95\%}$[0.44 ; 0.54] *** |
| | | 0.002 | x : p1 – 1 | x : 0.85 ± 0.023 | x : 0.518 $IC_{95\%}$[0.41 ; 0.61] *** |

| | | | | | |
|---|---|---|---|---|---|
| | | | 2-3 : p1 - 1 | 2-3 : 0.71 ± 0.018 | 2-3 : 0.546 $IC_{95\%}$[0.50 ; 0.59] *** |
| | | 0.005 | x : p1 – 1<br>2-3 : p1 - 1 | x : 0.82 ± 0.023<br>2-3 : 0.56 ± 0.016 | x : 0.546 $IC_{95\%}$[0.45 ; 0.63] ***<br>2-3 : 0.568 $IC_{95\%}$[0.52 ; 0.61] *** |
| | | 0.01 | x : p1 – 1<br>2-3 : p1 - 1 | x : 0.75 ± 0.021<br>2-3 : 0.42 ± 0.015 | x : 0.568 $IC_{95\%}$[0.47 ; 0.65] ***<br>2-3 : 0.595 $IC_{95\%}$[0.55 ; 0.63] *** |
| | | 0.02 | x : p1 – 1<br>2-3 : p1 - 1 | x : 0.67 ± 0.022<br>2-3 : 0.35 ± 0.014 | x : 0.581 $IC_{95\%}$[0.49 ; 0.66] ***<br>2-3 : 0.608 $IC_{95\%}$[0.57 ; 0.65] *** |
| | | 0.05 | x : p1 – 1<br>2-3 : p1 - 1 | x : 0.60 ± 0.021<br>2-3 : 0.25 ± 0.014 | x : 0.608 $IC_{95\%}$[0.52 ; 0.68] ***<br>2-3 : 0.615 $IC_{95\%}$[0.57 ; 0.65] *** |
| | | 0.1 | x : p1 – 1<br>2-3 : p1 - 1 | x : 0.52 ± 0.025<br>2-3 : 0.21 ± 0.014 | x : 0.636 $IC_{95\%}$[0.55 ; 0.71] ***<br>2-3 : 0.619 $IC_{95\%}$[0.58 ; 0.66] *** |
| | 30% - 70% | 0.001 | x : p1 – 1<br>2-3 : p1 - 1 | x : 0.95 ± 0.012<br>2-3 : 0.86 ± 0.011 | x : 0.475 $IC_{95\%}$[0.37 ; 0.57] ***<br>2-3 : 0.496 $IC_{95\%}$[0.45 ; 0.54] *** |
| | | 0.002 | x : p1 – 1<br>2-3 : p1 - 1 | x : 0.92 ± 0.012<br>2-3 : 0.76 ± 0.012 | x : 0.501 $IC_{95\%}$[0.39 ; 0.59] ***<br>2-3 : 0.540 $IC_{95\%}$[0.49 ; 0.58] *** |
| | | 0.005 | x : p1 – 1<br>2-3 : p1 - 1 | x : 0.84 ± 0.014<br>2-3 : 0.61 ± 0.013 | x : 0.540 $IC_{95\%}$[0.44 ; 0.63] ***<br>2-3 : 0.572 $IC_{95\%}$[0.53 ; 0.61] *** |
| | | 0.01 | x : p1 – 1<br>2-3 : p1 - 1 | x : 0.76 ± 0.015<br>2-3 : 0.45 ± 0.012 | x : 0.586 $IC_{95\%}$[0.49 ; 0.67] ***<br>2-3 : 0.597 $IC_{95\%}$[0.55 ; 0.64] *** |
| | | 0.02 | x : p1 – 1<br>2-3 : p1 - 1 | x : 0.67 ± 0.016<br>2-3 : 0.34 ± 0.011 | x : 0.610 $IC_{95\%}$[0.52 ; 0.69] ***<br>2-3 : 0.610 $IC_{95\%}$[0.57 ; 0.65] *** |
| | | 0.05 | x : p1 – 1<br>2-3 : p1 - 1 | x : 0.59 ± 0.019<br>2-3 : 0.24 ± 0.011 | x : 0.640 $IC_{95\%}$[0.56 ; 0.71] ***<br>2-3 : 0.613 $IC_{95\%}$[0.57 ; 0.65] *** |
| | | 0.1 | x : p1 – 1<br>2-3 : p1 - 1 | x : 0.90 ± 0.024<br>2-3 : 0.81 ± 0.019 | x : 0.651 $IC_{95\%}$[0.57 ; 0.72] ***<br>2-3 : 0.617 $IC_{95\%}$[0.58 ; 0.65] *** |
| | 45% - 55% | 0.001 | x : p1 – 0.986<br>2-3 : p1 – 0.997 | x : 0.98 ± 0.009<br>2-3 : 0.96 ± 0.009 | x : 0.213 $IC_{95\%}$[0.08 ; 0.33] *<br>2-3 : 0.493 $IC_{95\%}$[0.44 ; 0.54] *** |
| | | 0.002 | x : p1 – 0.993<br>2-3 : p1 – 0.998 | x : 0.99 ± 0.009<br>2-3 : 0.93 ± 0.011 | x : 0.231 $IC_{95\%}$[0.10 ; 0.35] **<br>2-3 : 0.504 $IC_{95\%}$[0.45 ; 0.55] *** |
| | | 0.005 | x : p1 – 0.989<br>2-3 : p1 – 0.998 | x : 0.94 ± 0.011<br>2-3 : 0.87 ± 0.014 | x : 0.523 $IC_{95\%}$[0.42 ; 0.61] ***<br>2-3 : 0.566 $IC_{95\%}$[0.52 ; 0.61] *** |
| | | 0.01 | x : p1 – 0.994<br>2-3 : p1 – 0.994 | x : 0.90 ± 0.014<br>2-3 : 0.75 ± 0.018 | x : 0.515 $IC_{95\%}$[0.41 ; 0.60] ***<br>2-3 : 0.555 $IC_{95\%}$[0.51 ; 0.60] *** |
| | | 0.02 | x : p1 – 0.993<br>2-3 : p1 – 0.995 | x : 0.83 ± 0.016<br>2-3 : 0.54 ± 0.020 | x : 0.584 $IC_{95\%}$[0.49 ; 0.66] ***<br>2-3 : 0.577 $IC_{95\%}$[0.53 ; 0.62] *** |
| | | 0.05 | x : p1 – 1<br>2-3 : p1 – 0.998 | x : 0.69 ± 0.020<br>2-3 : 0.29 ± 0.018 | x : 0.618 $IC_{95\%}$[0.53 ; 0.69] ***<br>2-3 : 0.607 $IC_{95\%}$[0.56 ; 0.64] *** |
| | | 0.1 | x : p1 – 1<br>2-3 : p1 - 1 | x : 0.53 ± 0.021<br>2-3 : 0.21 ± 0.012 | x : 0.633 $IC_{95\%}$[0.55 ; 0.70] ***<br>2-3 : 0.614 $IC_{95\%}$[0.57 ; 0.65] *** |
| Dominant by recessive (dominant major) | 10% - 90% | 0.001 | x : Recessive – 1<br>2-3 : Recessive - 1 | x : 0.89 ± 0.024<br>2-3 : 0.76 ± 0.018 | x : 0.399 $IC_{95\%}$[0.28 ; 0.50] **<br>2-3 : 0.518 $IC_{95\%}$[0.47 ; 0.56] *** |
| | | 0.002 | x : Recessive – 1<br>2-3 : Recessive - 1 | x : 0.86 ± 0.022<br>2-3 : 0.64 ± 0.017 | x : 0.529 $IC_{95\%}$[0.43 ; 0.62] ***<br>2-3 : 0.545 $IC_{95\%}$[0.50 ; 0.59] *** |
| | | 0.005 | x : Recessive – 1<br>2-3 : Recessive - 1 | x : 0.78 ± 0.023<br>2-3 : 0.49 ± 0.016 | x : 0.565 $IC_{95\%}$[0.46 ; 0.65] ***<br>2-3 : 0.581 $IC_{95\%}$[0.54 ; 0.62] *** |
| | | 0.01 | x : Recessive – 1<br>2-3 : Recessive - 1 | x : 0.74 ± 0.023<br>2-3 : 0.36 ± 0.014 | x : 0.593 $IC_{95\%}$[0.50 ; 0.67] ***<br>2-3 : 0.563 $IC_{95\%}$[0.56 ; 0.64] *** |
| | | 0.02 | x : Recessive – 1<br>2-3 : Recessive - 1 | x : 0.65 ± 0.023<br>2-3 : 0.30 ± 0.014 | x : 0.591 $IC_{95\%}$[0.50 ; 0.67] ***<br>2-3 : 0.609 $IC_{95\%}$[0.57 ; 0.65] *** |
| | | 0.05 | x : Recessive – 1<br>2-3 : Recessive - 1 | x : 0.58 ± 0.026<br>2-3 : 0.23 ± 0.014 | x : 0.616 $IC_{95\%}$[0.53 ; 0.69] ***<br>2-3 : 0.615 $IC_{95\%}$[0.57 ; 0.65] *** |

| | | | | | |
|---|---|---|---|---|---|
| | | 0.1 | x : Recessive – 1<br>2-3 : Recessive - 1 | x : 0.52 ± 0.026<br>2-3 : 0.19 ± 0.018 | x : 0.643 $IC_{95\%}$[0.56 ; 0.71] ***<br>2-3 : 0.616 $IC_{95\%}$[0.57 ; 0.65] *** |
| | 30% - 70% | 0.001 | x : Recessive – 1<br>2-3 : Recessive - 1 | x : 0.92 ± 0.012<br>2-3 : 0.67 ± 0.010 | x : 0.486 $IC_{95\%}$[0.38 ; 0.58] **<br>2-3 : 0.552 $IC_{95\%}$[0.51 ; 0.59] *** |
| | | 0.002 | x : Recessive – 1<br>2-3 : Recessive - 1 | x : 0.86 ± 0.012<br>2-3 : 0.54 ± 0.010 | x : 0.543 $IC_{95\%}$[0.44 ; 0.63] ***<br>2-3 : 0.574 $IC_{95\%}$[0.53 ; 0.62] *** |
| | | 0.005 | x : Recessive – 1<br>2-3 : Recessive - 1 | x : 0.76 ± 0.014<br>2-3 : 0.40 ± 0.009 | x : 0.574 $IC_{95\%}$[0.48 ; 0.66] ***<br>2-3 : 0.604 $IC_{95\%}$[0.56 ; 0.64] *** |
| | | 0.01 | x : Recessive – 1<br>2-3 : Recessive - 1 | x : 0.68 ± 0.015<br>2-3 : 0.32 ± 0.010 | x : 0.606 $IC_{95\%}$[0.52 ; 0.68] ***<br>2-3 : 0.613 $IC_{95\%}$[0.57 ; 0.65] *** |
| | | 0.02 | x : Recessive – 1<br>2-3 : Recessive - 1 | x : 0.59 ± 0.016<br>2-3 : 0.26 ± 0.009 | x : 0.626 $IC_{95\%}$[0.54 ; 0.70] ***<br>2-3 : 0.616 $IC_{95\%}$[0.57 ; 0.65] *** |
| | | 0.05 | x : Recessive – 1<br>2-3 : Recessive - 1 | x : 0.55 ± 0.020<br>2-3 : 0.21 ± 0.010 | x : 0.656 $IC_{95\%}$[0.57 ; 0.72] **<br>2-3 : 0.608 $IC_{95\%}$[0.57 ; 0.65] *** |
| | | 0.1 | x : Recessive – 1<br>2-3 : Recessive - 1 | x : 0.50 ± 0.022<br>2-3 : 0.21 ± 0.013 | x : 0.642 $IC_{95\%}$[0.56 ; 0.71] ***<br>2-3 : 0.591 $IC_{95\%}$[0.55 ; 0.63] *** |
| | 45% - 55% | 0.001 | x : Recessive – 0.995<br>2-3 : Recessive - 1 | x : 0.92 ± 0.010<br>2-3 : 0.67 ± 0.009 | x : 0.556 $IC_{95\%}$[0.46 ; 0.64] ***<br>2-3 : 0.543 $IC_{95\%}$[0.50 ; 0.59] *** |
| | | 0.002 | x : Recessive – 1<br>2-3 : Recessive - 1 | x : 0.87 ± 0.011<br>2-3 : 0.55 ± 0.009 | x : 0.544 $IC_{95\%}$[0.44 ; 0.63] ***<br>2-3 : 0.570 $IC_{95\%}$[0.53 ; 0.61] *** |
| | | 0.005 | x : Recessive – 1<br>2-3 : Recessive - 1 | x : 0.77 ± 0.013<br>2-3 : 0.37 ± 0.008 | x : 0.591 $IC_{95\%}$[0.50 ; 0.67] ***<br>2-3 : 0.607 $IC_{95\%}$[0.57 ; 0.64] *** |
| | | 0.01 | x : Recessive – 1<br>2-3 : Recessive - 1 | x : 0.68 ± 0.014<br>2-3 : 0.29 ± 0.008 | x : 0.623 $IC_{95\%}$[0.54 ; 0.70] ***<br>2-3 : 0.617 $IC_{95\%}$[0.58 ; 0.65] *** |
| | | 0.02 | x : Recessive – 1<br>2-3 : Recessive - 1 | x : 0.60 ± 0.016<br>2-3 : 0.25 ± 0.008 | x : 0.636 $IC_{95\%}$[0.55 ; 0.71] ***<br>2-3 : 0.610 $IC_{95\%}$[0.57 ; 0.65] *** |
| | | 0.05 | x : Recessive – 1<br>2-3 : Recessive - 1 | x : 0.54 ± 0.019<br>2-3 : 0.22 ± 0.010 | x : 0.657 $IC_{95\%}$[0.57 ; 0.72] ***<br>2-3 : 0.599 $IC_{95\%}$[0.56 ; 0.64] *** |
| | | 0.1 | x : Recessive – 1<br>2-3 : Recessive - 1 | x : 0.50 ± 0.022<br>2-3 : 0.20 ± 0.011 | x : 0.678 $IC_{95\%}$[0.60 ; 0.74] ***<br>2-3 : 0.585 $IC_{95\%}$[0.54 ; 0.62] *** |
| Dominant by recessive (recessive major) | 10% - 90% | 0.001 | x : Dominant – 1<br>2-3 : Dominant - 1 | x : 0.69 ± 0.025<br>2-3 : 0.37 ± 0.016 | x : 0.589 $IC_{95\%}$[0.50 ; 0.67] ***<br>2-3 : 0.589 $IC_{95\%}$[0.55 ; 0.63] *** |
| | | 0.002 | x : Dominant – 1<br>2-3 : Dominant - 1 | x : 0.61 ± 0.025<br>2-3 : 0.25 ± 0.014 | x : 0.617 $IC_{95\%}$[0.53 ; 0.69] ***<br>2-3 : 0.608 $IC_{95\%}$[0.57 ; 0.65] *** |
| | | 0.005 | x : Dominant – 1<br>2-3 : Dominant - 1 | x : 0.50 ± 0.028<br>2-3 : 0.17 ± 0.014 | x : 0.645 $IC_{95\%}$[0.56 ; 0.71] ***<br>2-3 : 0.606 $IC_{95\%}$[0.56 ; 0.64] *** |
| | | 0.01 | x : Dominant – 1<br>2-3 : Dominant - 1 | x : 0.44 ± 0.040<br>2-3 : 0.18 ± 0.020 | x : 0.642 $IC_{95\%}$[0.56 ; 0.71] ***<br>2-3 : 0.557 $IC_{95\%}$[0.51 ; 0.60] *** |
| | | 0.02 | x : Dominant – 1<br>2-3 : Dominant - 1 | x : 0.37 ± 0.067<br>2-3 : 0.33 ± 0.068 | x : 0.613 $IC_{95\%}$[0.52 ; 0.69] ***<br>2-3 : 0.491 $IC_{95\%}$[0.44 ; 0.54] *** |
| | | 0.05 | x : Dominant – 1<br>2-3 : Dominant - 1 | x : 0.50 ± 2.35<br>2-3 : 1 ± 4.946 | x : 0.409 $IC_{95\%}$[0.29 ; 0.51] **<br>2-3 : 0.364 $IC_{95\%}$[0.31 ; 0.42] *** |
| | | 0.1 | x : Dominant – 1<br>2-3 : Dominant - 1 | x : 1 ± 447.1<br>2-3 : 1 ± 1.718 | x : -0.008 $IC_{95\%}$[-0.14 ; 0.12] NS<br>2-3 : 0.240 $IC_{95\%}$[0.18 ; 0.30] ** |
| | 30% - 70% | 0.001 | x : Dominant – 1<br>2-3 : Dominant - 1 | x : 0.94 ± 0.016<br>2-3 : 1.06 ± 0.021 | x : 0.445 $IC_{95\%}$[0.33 ; 0.54] **<br>2-3 : -0.535 $IC_{95\%}$[-0.58 ; -0.49] *** |
| | | 0.002 | x : Dominant – 1<br>2-3 : Dominant - 1 | x : 0.92 ± 0.019<br>2-3 : 1.16 ± 0.029 | x : 0.469 $IC_{95\%}$[0.36 ; 0.56] **<br>2-3 : -0.509 $IC_{95\%}$[-0.55 ; -0.46] *** |
| | | 0.005 | x : Dominant – 1 | x : 0.97 ± 0.027 | x : -0.393 $IC_{95\%}$[-0.50 ; -0.28] ** |

| | | | | | |
|---|---|---|---|---|---|
| | | | 2-3 : Dominant – 0.993 | 2-3 : 1.33 ± 0.038 | 2-3 : -0.541 $IC_{95\%}$[-0.58 ; -0.49] *** |
| | | 0.01 | x : Dominant – 1<br>2-3 : Dominant – 0.649 | x : 1.08 ± 0.034<br>2-3 : 0.85 ± 0.043 | x : -0.584 $IC_{95\%}$[-0.66 ; -0.49] ***<br>2-3 : -0.578 $IC_{95\%}$[-0.62 ; -0.53] *** |
| | | 0.02 | x : Dominant – 0.97<br>2-3 : Recessive – 0.763 | x : 1.35 ± 0.040<br>2-3 : 0.28 ± 0.027 | x : -0.634 $IC_{95\%}$[-0.71 ; -0.55] ***<br>2-3 : 0.609 $IC_{95\%}$[0.57 ; 0.65] *** |
| | | 0.05 | x : Dominant – 0.555<br>2-3 : Recessive – 0.578 | x : 0.53 ± 0.043<br>2-3 : 0.22 ± 0.018 | x : -0.650 $IC_{95\%}$[-0.72 ; -0.57] ***<br>2-3 : 0.590 $IC_{95\%}$[0.55 ; 0.63] *** |
| | | 0.1 | x : Dominant – 0.982<br>2-3 : Dominant – 0.982 | x : 0.33 ± 0.270<br>2-3 : 0.79 ± 2.047 | x : 0.050 $IC_{95\%}$[-0.08 ; 0.18] NS<br>2-3 : 0.094 $IC_{95\%}$[0.03 ; 0.16] * |
| | 45% - 55% | 0.001 | x : Dominant – 0.938<br>2-3 : Recessive – 0.844 | x : 1.03 ± 0.010<br>2-3 : 0.72 ± 0.012 | x : -0.524 $IC_{95\%}$[-0.61 ; -0.42] ***<br>2-3 : 0.535 $IC_{95\%}$[0.49 ; 0.59] *** |
| | | 0.002 | x : Dominant – 0.76<br>2-3 : Recessive- 0.997 | x : 0.98 ± 0.013<br>2-3 : 0.58 ± 0.010 | x : -0.55 $IC_{95\%}$[-0.64 ; -0.45] ***<br>2-3 : 0.565 $IC_{95\%}$[0.52 ; 0.61] *** |
| | | 0.005 | x : Recessive – 0.782<br>2-3 : Recessive- 1 | x : 0.78 ± 0.013<br>2-3 : 0.38 ± 0.010 | x : 0.568 $IC_{95\%}$[0.47 ; 0.65] ***<br>2-3 : 0.601 $IC_{95\%}$[0.56 ; 0.64] *** |
| | | 0.01 | x : Recessive – 0.993<br>2-3 : Recessive - 1 | x : 0.69 ± 0.014<br>2-3 : 0.28 ± 0.008 | x : 0.620 $IC_{95\%}$[0.53 ; 0.69] ***<br>2-3 : 0.613 $IC_{95\%}$[0.57 ; 0.65] *** |
| | | 0.02 | x : Recessive – 1<br>2-3 : Recessive - 1 | x : 0.58 ± 0.016<br>2-3 : 0.24 ± 0.008 | x : 0.612 $IC_{95\%}$[0.55 ; 0.71] ***<br>2-3 : 0.613 $IC_{95\%}$[0.57 ; 0.65] *** |
| | | 0.05 | x : Recessive– 1<br>2-3 : Recessive - 1 | x : 0.54 ± 0.020<br>2-3 : 0.22 ± 0.009 | x : 0.661 $IC_{95\%}$[0.58 ; 0.73] ***<br>2-3 : 0.592 $IC_{95\%}$[0.55 ; 0.63] *** |
| | | 0.1 | x : Recessive – 1<br>2-3 : Recessive - 1 | x : 0.48 ± 0.022<br>2-3 : 0.20 ± 0.011 | x : 0.659 $IC_{95\%}$[0.58 ; 0.73] ***<br>2-3 : 0.57 $IC_{95\%}$[0.53 ; 0.61] *** |

Supplementary Table 3 : **Results of all the simulations under the model of pairwise incompatibility.** Each model is characterized by 3 parameters, the dominance (*d*) of the variants involved in the incompatibility, the admixture proportions in which population p1 and p2 are mixed to form a third population (*p*) and the mean of the exponential distribution from which is sampled the fitness impact of each incompatibility $(\bar{v})$. For each set of parameter, the minority ancestry after 1,600 generations on the sex chromosome and the autosomes as well as the proportions of model (over 1,000) in which this ancestry is minority. The median of the distribution of the ratio between the 50 lowest and highest recombining regions for all replicates as well as the 95% confidence interval of the value separately for the sex chromosome and the autosomes. Finally, the correlation coefficient between the average (across the 1,000 replicates) minor ancestry and the recombination rates for the X chromosome and the autosomes. The significance of the correlation is shown with different symbols, NS if the p-value is >0.05, * if 0.05>p-value>0.001, **if 0.001>p-value>2.2e-16 and *** if p-value < 2.2e-16.
